# Spatial Theta Cells in Competitive Burst Synchronization Networks: Reference Frames from Phase Codes

**DOI:** 10.1101/211458

**Authors:** Joseph D. Monaco, Hugh T. Blair, Kechen Zhang

## Abstract

Spatial cells of the hippocampal formation are embedded in networks of theta cells. The septal theta rhythm (6–10 Hz) organizes the spatial activity of place and grid cells in time, but it remains unclear how spatial reference points organize the temporal activity of theta cells in space. We study spatial theta cells in simulations and single-unit recordings from exploring rats to ask whether temporal phase codes may anchor spatial representations to the outside world. We theorize that an experience-independent mechanism for temporal coding may combine with burst synchronization to continuously calibrate self-motion to allocentric reference frames. Subcortical recordings revealed spatial theta cells with strong rate-phase correlations related to distinct theta phases. Simulations of bursting neurons and networks explained that relationship and, with competitive learning, demonstrated flexible spatial synchronization patterns when driven by low-dimensional spatial components from the recording data. Thus temporal coding synchrony may reconcile extrinsic and intrinsic neural codes.

## Introduction

Neural codes may map sensory or physical dimensions to intensity, like a rate code, or timing, like a phase code (cf. Brette, 2015). It remains unresolved how the apparent spatial metric revealed in grid cells (McNaughton et al., 2006; Moser & Moser, 2008) is computed in the brain as a rate-coding or a phase-coding process (Burak & Fiete, 2009; Zilli et al., 2009). Grid cells develop after place and border cells (Langston et al., 2010; Wills et al., 2010) and undergo experience-dependent distortion or control by geometric reference points in asymmetric or rotated environments (Krupic et al., 2015; Stensola et al., 2015; Savelli et al., 2017). These findings suggest the grid cell spatial metric is more local and malleable than universal and absolute. This characterization is at odds with the first-class grid responses formed by rate-coding continuous attractor models (Fuhs & Touretzky, 2006; Burak & Fiete, 2009) and the lack of a detailed mechanism for environmental resetting in phase-coding oscillatory interference models (N. Burgess, 2008; Hasselmo, 2008; Blair et al., 2008). Reconciling these gaps will elucidate the neural basis of spatial reference frames.

How can environmental cues reset a neural code for space? Path integration is the idiothetic process, analogous to angular integration for head direction (Zhang, 1996; Knierim et al., 1998), that guides the neural code for position using egocentric motion signals. Interacting with external sensory cues is critical for path integration to remain calibrated within a fixed spatial reference frame (Gothard et al., 1996; Etienne et al., 2004), but the nature of the interaction depends on the neural code. A path-integrating rate code may require direct activation of sensory associations with the cells coding for position (Widloski & Fiete, 2014). A path-integrating phase code may require temporal phase synchronization with a sensory feedback signal (Monaco et al., 2011; Blair et al., 2014). Phase-code models (N. Burgess, 2008; Hasselmo, 2008) have previously implemented resetting as instantly forcing oscillators to zero phase, similar to experimental stimulation- or task-evoked theta reset (Buzsáki et al., 1979; Williams & Givens, 2003). Rather than abrupt resets, the spatial calibration of grid cells may be predominantly mediated by boundaries (Hardcastle et al., 2015) in a way that reflects continuous gating between extrinsic and intrinsic information streams (Carpenter et al., 2015; Savelli et al., 2017). It is unclear how place-to-grid feedback may support this form of calibration in attractor network or phase synchronization models.

Which pathway would place-to-grid feedback take? The hippocampal formation is itself a loop-like structure, but subregions CA1/CA3 also form bidirectional loops with subcortical structures that regulate the septo-hippocampal theta rhythm (6–10 Hz; Leranth et al., 1999; Ruan et al., 2017). Theta-rhythmic activity propagates through the circuits of the septum, mammillary bodies, and anterior thalamus via excitatory burst synchronization (Vertes et al., 2001; Tsanov et al., 2011; Welday et al., 2011). Bursting aids neurocomputation and signal transmission by overcoming synaptic failure, facilitating transmitter release, selecting resonant inputs, and/or evoking synaptic plasticity (Lisman, 1997; Izhikevich, 2007). In this paper, we study the hypothesis that theta bursting and spatial inputs create a spatial phase code that supports flexible learning of spatial synchronization patterns. We theorize that this spatial phase code is reflected in correlations between firing rate and phase. We recorded theta cells from a constellation of hippocampal and subcortical areas in freely exploring rats to look for spatial phase information and rate-phase correlations. We modeled intrinsic theta bursting in oscillatory neuronal network models to demonstrate 1D and 2D phase-code synchronization to, respectively, artificial and path-integration-like phase codes. We discuss the results with respect to the hippocampo-entorhinal spatial metric, but the rate-to-phase calibration mechanism may subserve other brain systems.

## Results

Our approach combines mechanistic models of burst synchronization with information theoretic and statistical modeling analyses of theta cell recordings. First, we present 1D simulations of spatial bursting cells that test how they entrain a nonspatial target neuron to a spatial phase code. Second, we present open-field recordings of theta cells in rats to quantify spatial phase coding and study a statistical model to isolate trajectory-based confounds of spatial activity. Third, we construct a 2D data-driven generative model of spatial inputs for competitive bursting network simulations that characterize dynamical constraints on environmental phase attractors.

### Spatial ‘phaser’ bursting models lock to distinct theta phases

To model spatial theta cells, we defined a two-variable, nonlinear integrate-and-fire model of intrinsic bursting, meaning that the bursting derives from internal dynamics and not external fluctuations. This bursting model (Methods) is a variation on Izhikevich’s hippocampal low-threshold burster (Izhikevich, 2007, p.310), which can fire single spikes or bursts of varying intensity depending on input strength. Its dynamics implement burst termination with adaptive feedback analogous to the slow calcium- or voltage-gated activation of outward currents (*I*_*AHP*_ or *I*_*K*(*Ca*)_) observed in hippocampal and midbrain bursting neurons (Traub et al., 1991; Amini et al., 1999). For recording phase, spiking simulations (Methods) tracked a reference theta wave at *f*_*θ*_ = 7.5 *s*^−1^, the typical burst rate of our theta-cell data below.

Mehta *et al* (2002) posited that combining inhibitory theta input with excitatory ramping input is a robust mechanism for creating a temporal code from a rate code: Lower excitation delays firing to the periodic inhibitory minimum (theta trough) and higher excitation advances firing until maximum inhibition (theta peak). This precession of activity conveys information about the rate-coded input in the theta phase of the output (Discussion).

To test temporal coding in the theta-bursting model, we implemented the Mehta mechanism by combining theta inhibition with excitation from a ramping input function *F*_ramp_ (equation (4); Methods). With certain bursting (Table 2) and gain (Table 3) parameters, we call this a ‘phaser’ cell. We demonstrate a phaser cell simulation (Figure 1A) in which *F*_ramp_ is a triangle wave (green). For low excitatory input, the phaser (Figure 1A, blue trace and spike raster) emits single spikes near theta peak (zero phase) every few theta cycles (gold highlights, *Low1* and *Low2*). For high excitatory input, the phaser bursts with spike triplets near theta trough (−*π*/*π* phase) every other theta cycle (gold highlight, *High*). This cycle-skipping rhythmicity is consistent with observations in medial entorhinal cortex and the head direction system (Deshmukh et al., 2010; Brandon et al., 2013; Discussion). Expanded intervals (Figure 1B) reveal the range of burst modulation (blue traces) and the shift in timing to earlier phases (middle) relative to the reference theta wave (magenta). More frequent bursts at earlier theta phases suggest the negative correlation between rate and phase entailed by the Mehta mechanism (2002). To quantify this correlation, we sampled spiking for a longer triangle wave with a varying cycle period. Phase distributions and the rate-phase correlation (*n* = 399 nonzero input bins out of 512; *r* = –0.809; Figure 1D) show clear precession from peak to trough (0 to −*π*) across average firing rate. Thus this ‘negative’ phaser forms a Mehta-like phase code of its input function that will most strongly entrain a post-synaptic target to the trough of the theta rhythm.

**Figure 1.**
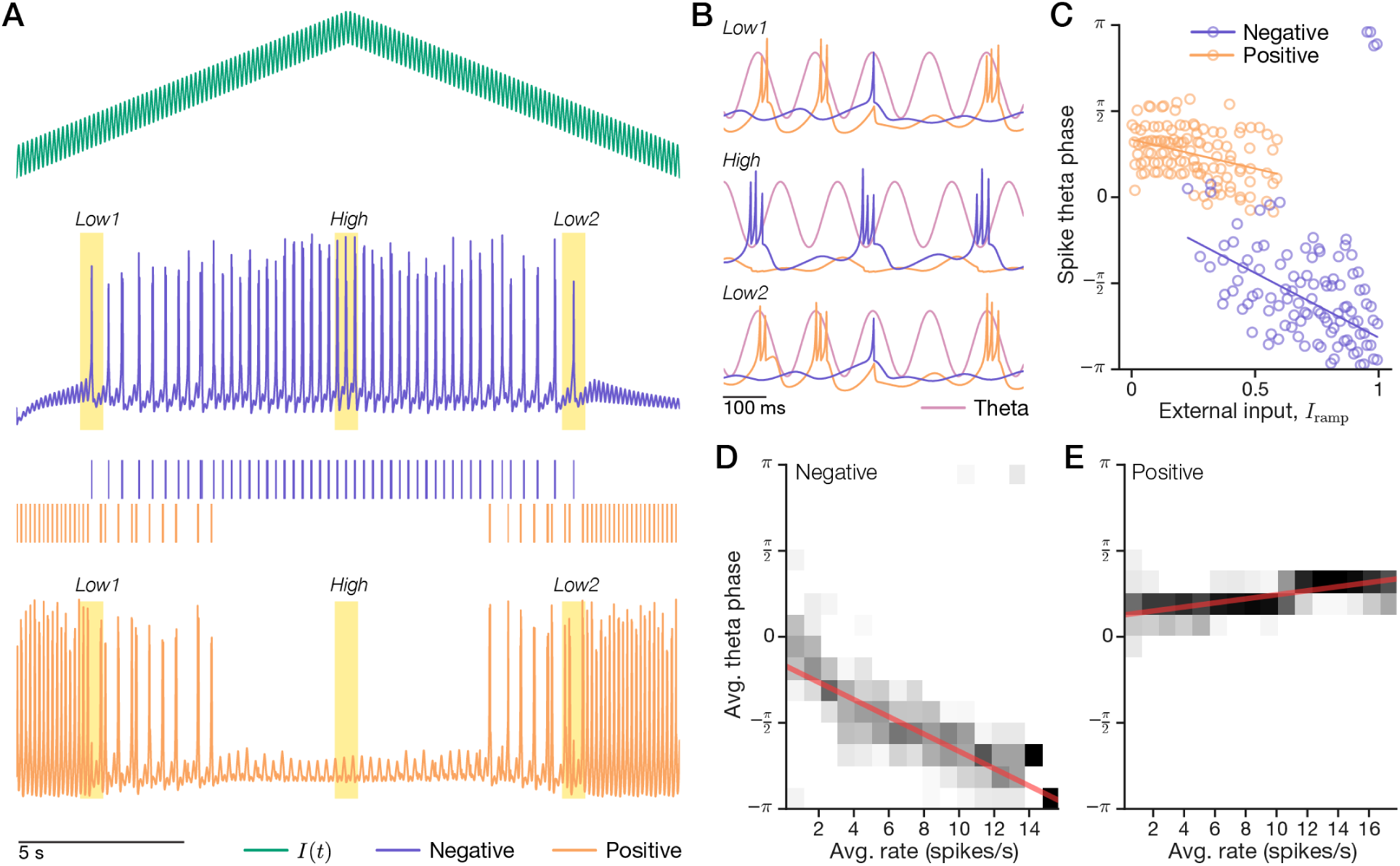
Excitatory input to theta-bursting neuron models creates rate-phase correlations. A bursting model (blue, ‘negative’) with combined external and theta input (green) suppresses another bursting model (orange, ‘positive’). (A-C) A 20-s simulation. (A) A triangle-wave input (top) produced a range of spiking (*Low1*, *Low2*) and bursting (*High*) in the negative cell (middle) and a complementary pattern in the positive cell (bottom). (B) Expanded intervals show the reference theta wave (magenta). (C) A scatter plot shows spike theta-phase across input levels. Stronger inputs caused earlier firing (phase precession) in the negative cell and silenced the positive cell. Lines, circular-linear regressions. (D+E) A 1-hr simulation of 10-s to 62-s triangle-wave cycles sampled average firing rates and phases. Rate-phase correlations (grayscale, phase distributions conditioned on rate) revealed that input level comodulated rate and phase. For higher firing rates, the negative cell strongly precessed to earlier phases (D) but the positive cell processed more weakly to later phases (E). Red lines, circular-linear regressions.

How can a minimal bursting model entrain more than one phase? We propose a parsimonious mechanism allowing for simultaneous entrainment to theta peak: A theta cell with strong excitatory theta input whose activity is suppressed by a negative phaser. To avoid the additional degrees-of-freedom and parameter tuning for an interneuronal subnetwork, we modeled the lateral inhibition as feedforward inhibition with a slow 100-ms conductance (equation (5); Table 3). The ‘positive’ phaser (Figure 1A, orange trace and spike raster) bursts in-phase with theta when disinhibited by weak input to the negative phaser (highlights, *Low1* and *Low2*; Figure 1B, top and bottom). The negative and positive phasers fire in complementary patterns as the input changes across the simulation (Figure 1A). The positive phaser appears to precess with stronger inputs (Figure 1C), but the suppressive inhibition means that its firing rate is increasing as the external input goes to zero. Thus the rate-phase correlation mediated by the input is positive (*n* = 351 nonzero input bins out of 512; *r* = 0.705; Figure 1E). This positive phaser procession is shallower than the negative phaser precession (Figure 1D) that is directly driven by the external input. Thus a simple connectivity pattern between theta cells may permit multiplexed entrainment to the peak and trough of the theta rhythm.

### Competitive learning synchronizes a 1D spatial phase code

Can these phaser cells create spatial synchronization patterns? We constructed a 1D spatial model from 64 spatial tuning functions, each representing a particular location similar to a place field. However, if these were the only spatial inputs, then the complementary firing patterns of negative and positive phasers (Figure 1A) would entail that positive phasers only have long-range, not local, spatial responses. To equalize the diversity of spatial responses between negative and positive phasers, we added 64 ‘inverse’ spatial functions representing the long-range complements of the local tuning functions (Figure 2A, purples; Methods). Each of these spatial inputs drives one of 128 negative phasers (Figure 2A, blues) connected (equation (5)) to one of 128 positive phasers (oranges). Example (*x* = 0.5) joint space-phase distributions for the 4 phaser/tuning subtypes (Figure 2B) show the resulting spatiotemporal patterns available for synchronization (for the entire network, see Supplementary Figures 1–4).

**Figure 2.**
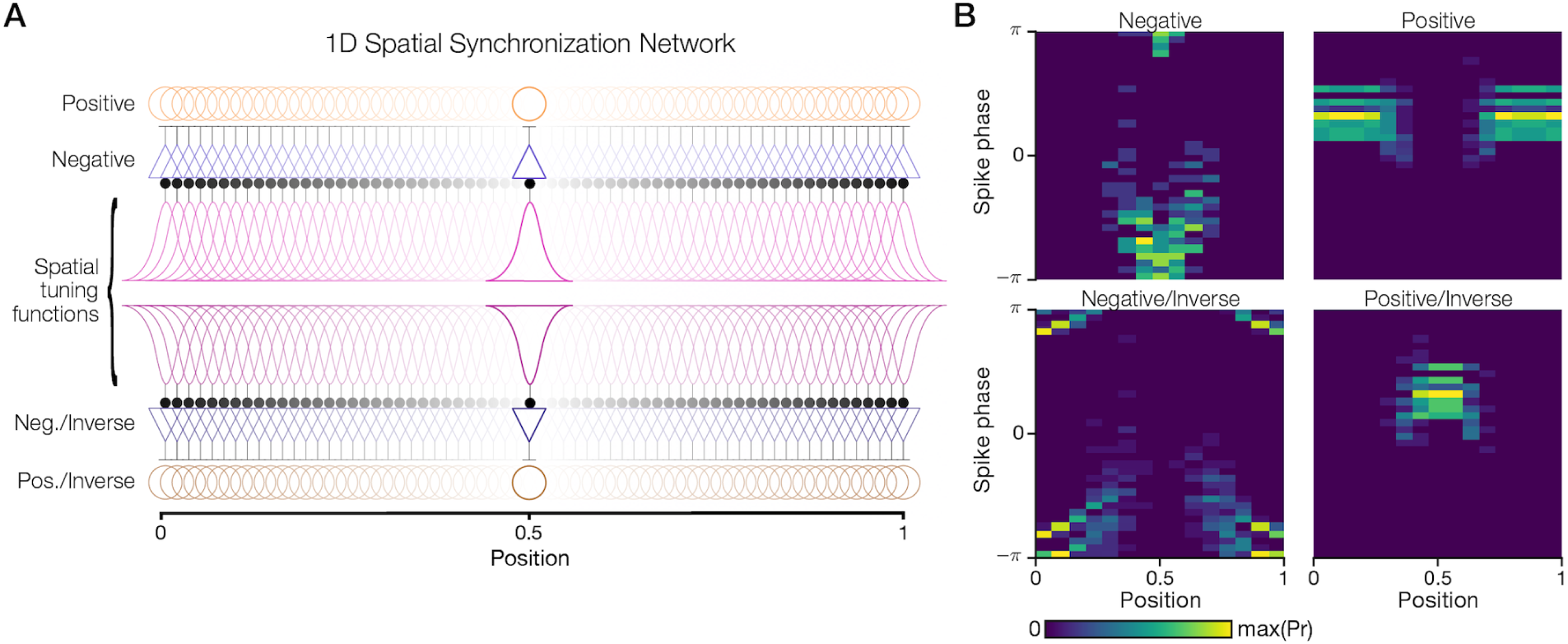
A 1D spatial network creates a palette of space-phase distributions. A set of 64 local tuning functions for a 1D space on {0, 1} and their corresponding inverse (long-range) tuning functions drive 128 pairs of negative/positive model phasers. (A) The tuning functions (purples: upper/local, lower/inverse) evenly cover the space and excite (filled circles) the negative phasers (blues). The negative phasers in turn suppress (T-bars) their paired positive phasers (oranges). The subnetwork at 0.5 is highlighted. (B) A 1-hr simulation sampled spike phase for a 1-min triangle-wave trajectory spanning the space. For the highlighted phasers in (A), the resulting joint space-phase distributions of spike timing create distinct spatiotemporal patterns around the theta trough (−*π*/*π* phase) and the falling phase ({0*, π*/2}).

Can this phaser activity entrain a 1D spatial phase code? We devised a binary phase-code target consisting of an anti-phase fixed point near *x* = 0 and an in-phase fixed point near *x* = 1 (Figure 3A). This pattern associates the opposing ends of the space with opposing phases of the theta cycle. We computed the vector cosine similarity between the phasers’ joint space-phase distributions and the phase-code target as a basis for competitively selecting active synapses (Methods). This winners-take-all (WTA; Table 4) method competed local against long-range (inverse) phasers within the negative/positive phaser subtypes. The resulting weights (Figure 3B) show the anti-phase fixed-point supported by negative/local phasers (left) and the in-phase fixed-point supported by positive/inverse phasers (right). To visualize the trained network, a weighted average of the phaser distributions (Figure 3C) revealed a qualitative match to the phase-code target in which the anti-phase mode (right) was more sharply defined than the in-phase mode (left). This pattern suggests spatial synchronization with phasers is possible.

**Figure 3.**
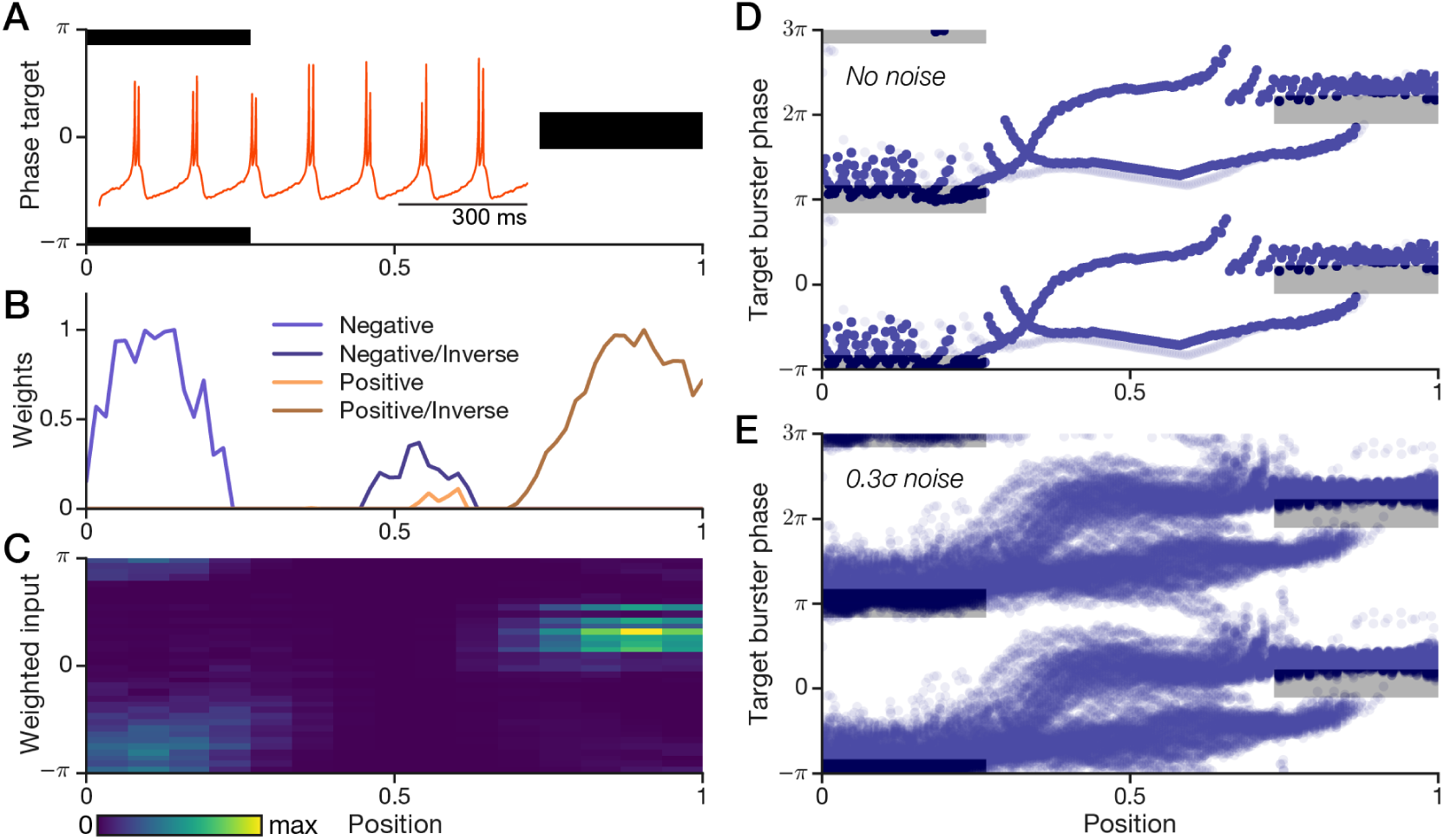
Competitively trained phasers synchronize a theta cell to a phase code. We devised an artificial spatial phase code and a nonspatial theta-cell model; competitive weights associating the 1D phaser network (Figure 2A) to the phase code allowed phaser input to spatially synchronize the theta cell. (A) The phase code target (black) has an anti-phase mode (left, −*π*/*π*) and an in-phase mode (right, zero phase). The target bursting cell (inset) random-walks across theta phase (Supplementary Figure 5). (B) Competitive 20%-winners-take-all weights (Methods). (C) Weighted average of the joint space-phase distributions of the phaser network. (D+E) Hour-long simulations of the target theta-cell burst timing on a 1-min triangle-wave trajectory between 0 and 1. The theta cell was simulated without (D) and with (E) intrinsic phase noise. Multiple theta cycles are shown (*y*-axis) for clarity.

To test phaser synchronization of downstream neuron, we created an intrinsic theta-bursting cell with simplified bursting dynamics (equation (6); Table 4; Methods) that emits doublets without cycle skipping. The intrinsic burst rate was approximately tuned to the reference theta frequency *f*_*θ*_. Without phaser input, this ‘target burster’ (Figure 3A, inset) randomly drifts across theta phase with a noise value (*σ*, Table 4) that randomizes its phase across a 30-s simulation (Supplementary Figure 5). To verify synchronization to the phase-code target, we simulated the phaser-target network with triangle-wave trajectories (1-min period, 1-hr duration). Plots of burst timing for simulations (Methods) without noise (Figure 3D) and with noise (Figure 3E) reveal stereotyped phase trajectories locked to both the in-phase and anti-phase fixed points (gray rectangles). The upper branch of the synchronization pattern, moving left toward *x* = 0, smoothly precesses to the earlier anti-phase fixed point; the lower branch, moving right toward *x* = 1, slowly processes to the fixed point until jumping discontinuously ahead of it (Figure 3D+E). The height of the burst-timing channels on either side, approximately a quarter cycle (Figure 3D+E), indicate the degree of phase misalignment tolerated in the target burster. While this tolerance shows the phaser synchronization does not act perfectly, it robustly prevents substantial drift from the phase-code target. This pattern holds across a range of noise levels and input gains (Supplementary Figure 6). Thus a spatial network of phaser cells can robustly synchronize a noisy theta-bursting neuron to an artificial spatial phase code.

### Theta cell recordings reveal spatial phaser patterns

To study space-phase representations in biological theta cells, we recorded single-unit data from rats foraging an open 80-cm cylindrical arena with tetrodes in theta-rhythmic sites including septum, hippocampus, thalamus, midbrain, and other subcortical areas (Methods). Recording sessions were longer than typical spatial navigation experiments (*n* = 110; mean, 2.1 hours; range, [0.76, 3.28]) to sufficiently sample phase differences across the environment. In all, 671 uniquely identified theta cells were recorded in 8 rats, resulting in 1,073 cell-session recordings for analysis.

Some theta cells were clearly modulated by space. An example spatial theta cell from the lateral septum fired preferentially on the west/southwest half of the arena (Figure 4A) with a peak firing-rate around 12 spikes/s (Figure 4B) on an adaptively-scaled spatial map of average firing rate (‘ratemap’; Methods). To verify this as a theta cell, its temporal autocorrelation (Figure 4C) revealed a theta rhythmicity index of 0.392 and its phase distribution relative to the hippocampal local field potential (LFP) theta rhythm (Figure 4D) revealed a theta modulation index of 0.288 (Methods) and a preference for anti-phase (−*π*/*π*) activity. However, the spatial map of average firing phase (‘phasemap’; Figure 4E) shows that the cell preferred in-phase firing (greens) in low firing-rate regions, and anti-phase (pinks) in high firing-rate regions. We computed ‘coherence maps’ by darkening phasemap pixels by phase variance (Methods). The coherence map (Figure 4F) shows that the spatial phase pattern holds in the arena center, but loses coherence along the west wall. Example intervals of spikes and LFP theta-phase show the transition from single spikes to bursts between periods of low and high firing rates (Figure 4G). Does this spatial theta cell carry correlations between rate and phase, as predicted by the Mehta mechanism? Indeed, the rate-phase correlation (Figure 4H) reveals a strong negative relationship (circular-linear regression: *n* = 3,190 pixels, *r* = –0.836; Methods) similar to our negative phaser model (Figure 1C, blue). Thus some theta cells may convert spatial inputs into spatial phase codes.

**Figure 4.**
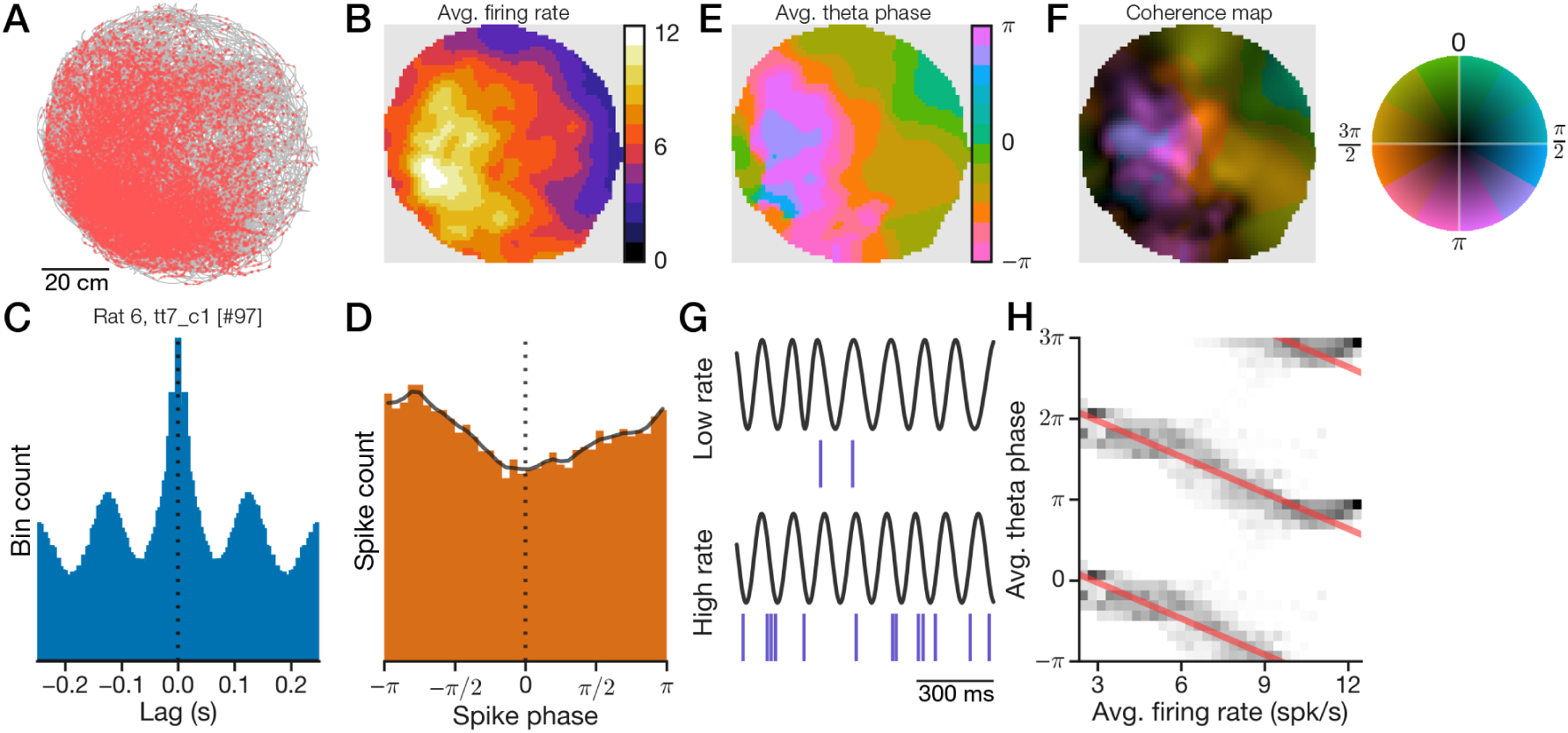
An example theta cell has spatial rate-phase correlations. Recordings from subcortical theta cells were made in rats exploring an 80-cm cylinder for long durations. (A) Spikes (red dots) from a lateral septal cell demonstrated spatial selectivity for one side of the arena (gray line, trajectory). (B) Adaptively smoothed map (Methods) of average firing rate. (C) Spike autocorrelogram (for computing theta rhythmicity index; Methods). (D) Distribution of spike phase relative to hippocampal theta (gray line, 10° moving average for computing theta modulation index; Methods). (E) Map of average theta-phase of firing. (F) Average phases from (E) composited with a saturation mask representing maximum-normalized phase coherence. Inset, phase-coherence color wheel. (G) Sample 1-s traces of LFP theta rhythm and spikes for low/high (top/bottom) periods of firing. (H) Conditional phase distributions along rate (grayscale), based on average rate (B) and phase (E) pixels, with circular-linear regression lines (red). Multiple theta cycles shown for clarity.

Are rate-phase correlations characteristic of spatial phase codes in theta cells? We computed spatial phase information *I*_phase_ with critical value *α* = 0.02 (Methods). Theta cell recordings with statistically significant *I*_phase_ (*n* = 233) have peak firing rate (median, 7.35 spikes/s), estimated burst frequency (7.66 *±* 0.44 *s*^−1^), and theta rhythmicity index (median, 0.365) values comparable to those of nonspatial cells (Supplementary Figure 7). Spatial recordings demonstrated a wide range of *I*_phase_ values (median, 0.36 bits; range, [0.012, 3.67]; Supplementary Figure 8A). How does spatial phase information relate to rate-phase correlations? Based on circular-linear regressions for each cell (like Figure 4H, red lines), we estimated the total phase shift for each recording (Methods). Total phase shifts for nonspatial recordings (Figure 5A, contours) were distributed around zero regardless of *I*_phase_, whereas total phase shifts for spatial recordings (blue circles) were strongly negative or positive even for low *I*_phase_ values. Both rate-phase correlation coefficients and total phase shifts were more broadly distributed for spatial cells than nonspatial cells (Supplementary Figure 8). Thus spatial theta cells may carry spatial phase information via negative and positive rate-phase correlations.

**Figure 5.**
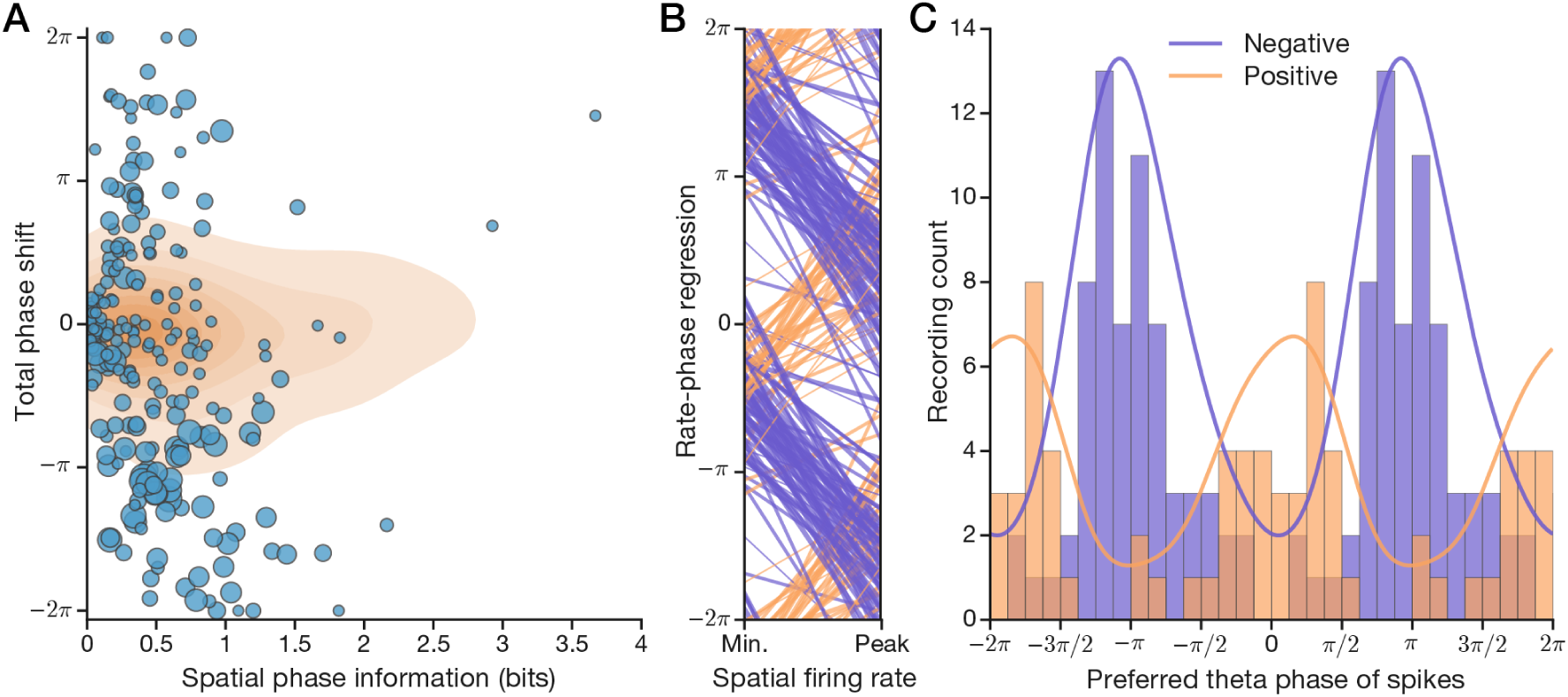
Biological phasers split into theta-segregated negative/positive subtypes. (A) To find phaser-like recordings, we compared spatial phase information with the total phase shift estimated from rate-phase correlations of all theta cells. Background contours show the distribution of 840 nonspatial theta cell recordings. Circles show 233 spatial recordings; circle size increases with correlation strength. (B+C) Phaser criteria (Results) selected 101 spatial recordings. Multiple theta cycles shown for clarity. (B) Rate-normalized rate-phase regression lines for negative (blue) and positive (orange) biological phaser recordings. (C) Preferred theta phases for negative/positive phasers are shown as histograms (positive composited over negative) and kernel-density estimates (lines, *π*/4 bandwidth Gaussian). On average, negative/positive phasers prefer anti-phase/in-phase firing.

What do these spatial phase-coding theta cells look like? Let’s define criteria for ‘biological phaser’ cells: peak firing rate ≥ 3.5 spikes/s, significant *I*_phase_ ≥ 0.1 bits, significant rate-phase correlation *|r| >* 0.2, and absolute phase shift ≥π/4. These criteria select 101/233 spatial recordings from 5/8 rats (Supplementary Figure 9). Like the model phasers, biological phasers had functional subtypes based on whether they fired earlier (negative, *n* = 65) or later (positive, *n* = 36) at higher firing rates. To evaluate their temporal organization, the rate-phase regression lines for each cell (Figure 5B) show that negative (blue) and positive (orange) phasers started firing during the rising phase of theta ({−π, 0}) and then, with increasing firing rate, precessed or processed on opposing paths to the falling phase ({0, π}). This rate-modulated phase pattern spans the theta cycle, but on average the negative phasers preferred trough firing and the positive phasers preferred peak firing (Figure 5C) as predicted by the model phasers (Figure 1C). Phaser cells constituted 13.2% of recordings from the septum (Table 1) where they were found predominantly in the dorsal/intermediate aspects of lateral septum (Supplementary Figure 10). For negative phaser cells (Figure 6A-E), ratemaps (top row) revealed diverse spatial representations including place-like fields (A+B), broad gradient-like fields (C+D, showing remarkably similar responses from different rats), and boundary responses along the arena wall (E). High firing rates (Figure 6A-E, top row) generally corresponded to pre-trough timing (middle row, blues/pinks). Many conditional phase distributions (Figure 6A-E, bottom row, grayscale) show that precession halted once the cell precessed past theta trough; note that this nonlinearity means that some regression lines (Figure 5B) overestimate phase shifts. Positive phaser cells (Figure 6F-J) likewise revealed diverse spatial modulation, but the responses were more subtle, involving higher baseline firing rates (F-I, top row) and heterogeneous compositions of boundary-like and place-like selectivity. Positive phasemaps (Figure 6F+H-J, middle row) showed shifts from pre-theta-peak (greens) to post-theta-peak (blues) that were evident in shallow rate-phase regressions centered on zero phase (bottom row); recording 52 (G) was an oddball with procession centered on theta trough. To quantify spatial differences between negative and positive phasers, the widely-used Skaggs information measure (1993; Methods) corroborated that negative phaser spikes carried more spatial content (negative: *n* = 47 unique cells; 0.381 *±* 0.06 bits/spike, mean *±* s.e.m.; positive: *n* = 24, 0.111 *±* 0.048; *post hoc* log-transformed Welch’s *t* = −3.92, *p* = 0.0002). This difference is consistent with our model (Figure 2A) where only the negative phasers are driven directly by spatial inputs. Thus biological phaser cells, prominently in lateral septum, represent diverse spatiotemporal relationships consistent with the phaser model.

**Figure 6.**
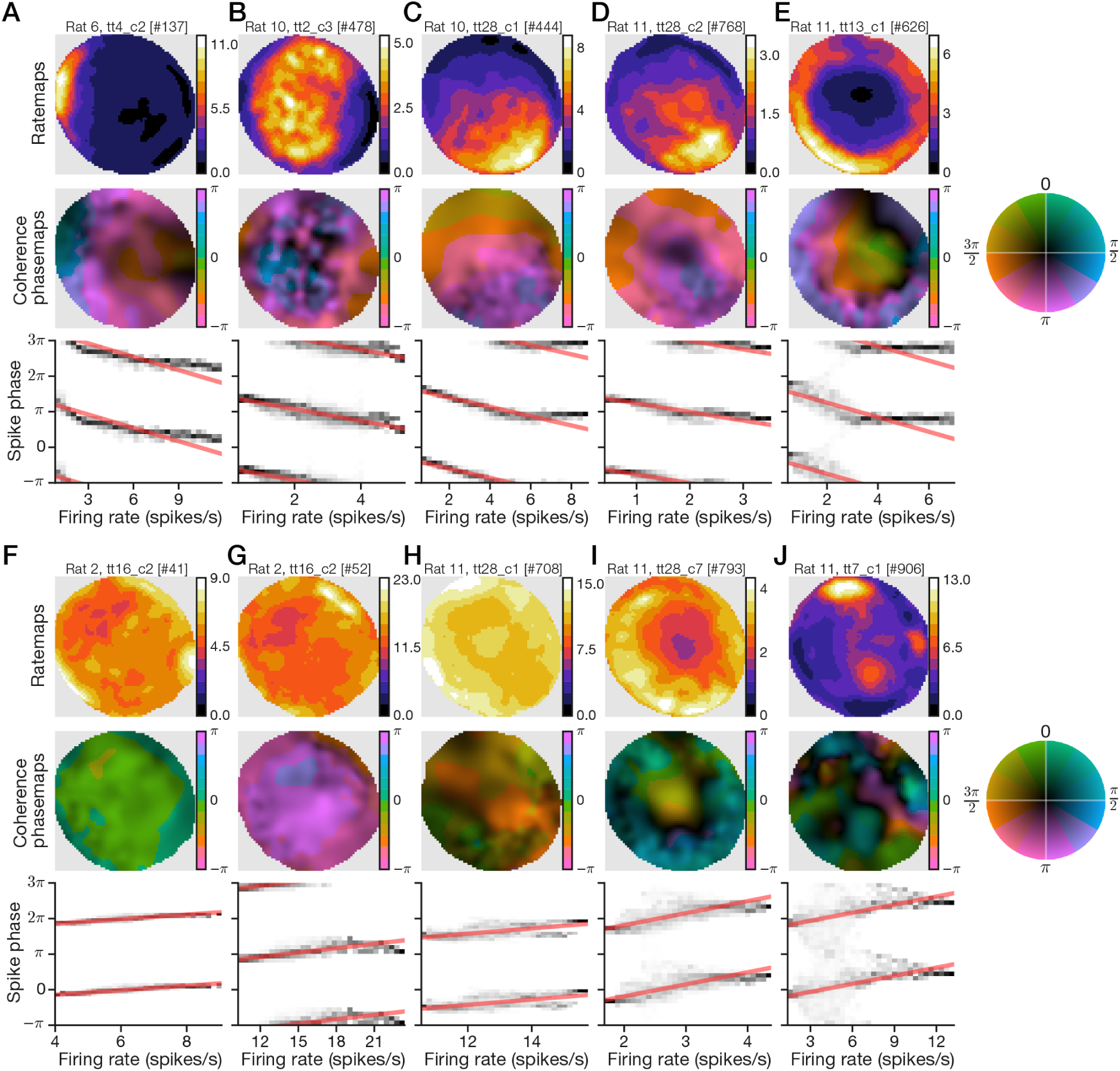
Example phaser recordings reveal diversity of spatial phase codes. For 5 example recordings of negative (A-E) and positive (F-J) biological phasers, we show the ratemap (top), phase-coherence map (middle), and conditional phase distributions with rate-phase regression lines (bottom, like Figure 4H). Note that peak firing rates (A-J, top, colorbar axes) are consistent with the restricted range of biological phaser firing rates (Supplementary Figure 9). Negative phasers generally showed stronger spatial modulation and rate-phase correlations than positive phasers (Results).

**Table 1.**
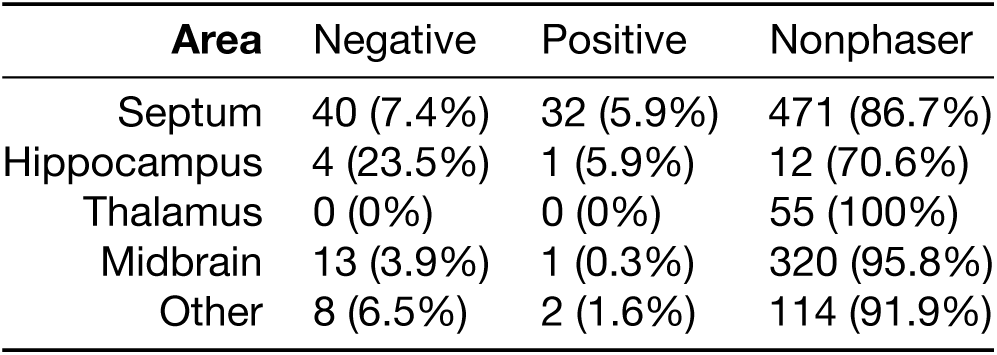
Counts of theta cell recordings by brain area and phaser subtype.

| Area | Negative | Positive | Nonphaser |
| --- | --- | --- | --- |
| Septum | 40 (7.4%) | 32 (5.9%) | 471 (86.7%) |
| Hippocampus | 4 (23.5%) | 1 (5.9%) | 12 (70.6%) |
| Thalamus | 0 (0%) | 0 (0%) | 55 (100%) |
| Midbrain | 13 (3.9%) | 1 (0.3%) | 320 (95.8%) |
| Other | 8 (6.5%) | 2 (1.6%) | 114 (91.9%) |

Do biological phaser recordings reflect exclusively spatial inputs? To quantify contributions from trajectory-based factors, we computed spike information and modulation indexes for direction and speed (Methods). As a spatial baseline, regressing Skaggs information onto spatial phase information *I*_phase_ yielded slope 0.831 (Supplementary Figure 11A), indicating that spike phase contributes ∼20.4% to the information rate beyond spike position alone. In contrast, regressing the spike information content for direction or speed onto *I*_phase_ yielded 0.103 and 0.036, respectively (Supplementary Figure 11B+C), indicating minimal coding overlap between *I*_phase_ and direction or speed. However, modulation indexes based on deflection of firing rates across direction (*n* = 101 phaser recordings; median, 0.379) and speed (0.318) suggested dependence of phaser cell activity on the trajectory (Supplementary Figure 11D+E). Thus phaser firing rate, but not phase, may reflect spatial-behavioral confounds which must be resolved.

### Statistical model of spatial drive isolates inhomogeneous directionality

The main behavioral confound is trajectory-biased sampling of cells whose directionality may vary by location. Spurious spatial activity may result from directionally biased visits to a particular location by the animal for which the recorded cell happened to have a similar directional preference. The problem is exacerbated by inhomogeneously directional cells that may exhibit a range of directional preferences across the environment. For example, a cell responding to anti-clockwise movement during running may produce a boundary-like response along the wall if the animal only runs anti-clockwise along the wall. To evaluate this confound, we studied a Poisson-distributed generalized linear model (GLM) of spatial and trajectory variables. GLMs have been shown to learn independent spatial and directional contributions to firing that avoid trajectory-based biases (Acharya et al., 2016). We fit the GLM independently to every cell recording for each element of a 3 × 3 spatial grid spanning the arena (Methods) to capture inhomogeneous changes in spatial or directional selectivity. The model is trained to predict spike counts across 300-ms intervals *i*

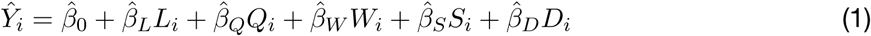

where *L* and *Q* are linear and quadratic spatial variables, *W* is a sigmoidal wall-proximity signal, *S* is linear speed, and *D* is movement direction. *L*, *Q*, and *W* are purely spatial whereas *S* and *D* capture the animal’s velocity vector, so we call this the LQW-SD model. The spatial predictors are more reliable over the training intervals than the velocity predictors. To address this asymmetry, we trained LQW-SD on standardized predictors as a ridge regression with *ℓ*_2_-regularization (Hastie et al., 2009). To maximally expose inhomogeneous directionality, we chose the regularization penalty that optimized the trade-off between maximizing the directional component of the model and minimizing the spike prediction error (Supplementary Figure 12; Methods). To quantify inhomogeneity, we computed a directional coherence index (DCI) on {0, 1} measuring alignment of the 9 *β*_*D*_ vectors across the grid; to quantify directionality, we computed a directional strength index (DSI) on {0, 1} measuring the magnitude of *β*_*D*_ relative to the other predictors (Methods). DCI for phasers (*n* = 69 unique cells; median, 0.315) showed higher coherence than nonphasers (*n* = 631; 0.213; *post hoc* Mann-Whitney *U* = 15,567, *p* = 0.0001). DSI for phasers (median, 0.0187) and nonphasers (0.0133) found similarly low directionality (*U* = 18,258, *p* = 0.0277) but nonphasers were more widely distributed (range, [0, 0.199]) than phasers ([0.002, 0.094]; Supplementary Figure 13). Biological phaser cells thus exclude both coherent (high DCI, high DSI) and inhomogeneous (low DCI, high DSI) directionality.

What does LQW-SD reveal about spatial predictors? Like DSI for directionality, we computed the relative strength of each model variable (equation (7); Methods). Box plots (Figure 7A) show the distribution of variable weights for phasers (green; *n* = 69 unique cells) and nonphasers (gray; *n* = 631). Both cell types had similar central tendencies with nonphasers exhibiting wider ranges of variable strengths. Spatial factors overwhelm the wall and trajectory variables, such that *L* and *Q* constitute approximately 30% and 60% of the model weight, respectively. Wall/boundary cells are (by observation) a small number within the dataset, but are the *S* and *D* factors really so low? We standardized predictors for training, but the trajectory-based signals may be highly non-normal. In that case, the importance of a model variable should be measured instead by its effective range of contribution to predictions. For each variable *X*, we computed its contribution

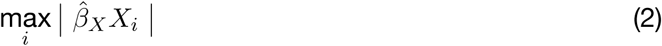

across time intervals *i* and sum-normalized the variables (Methods). The contribution profile (Figure 7B) was also dominated by *L* and *Q*, but *W*, *S*, and *D* contributions were enhanced relative to the strength profile (Figure 7A). Wall and direction variables constituted approximately 8% each of the total contribution and nonphasers revealed a wide range of speed contributions (Figure 7B, *S*, gray) consistent with extensive speed modulation in space-related brain areas (Fuhrmann et al., 2015; Kropff et al., 2015). Sorted matrixes of cell-level data confirmed this pattern and showed an inverse relationship between spatial and speed contributions (Supplementary Figure 14). Thus LQW-SD revealed that spatial factors trade off with speed modulation in theta cells and that phasers were overwhelmingly spatial, not directional.

**Figure 7.**
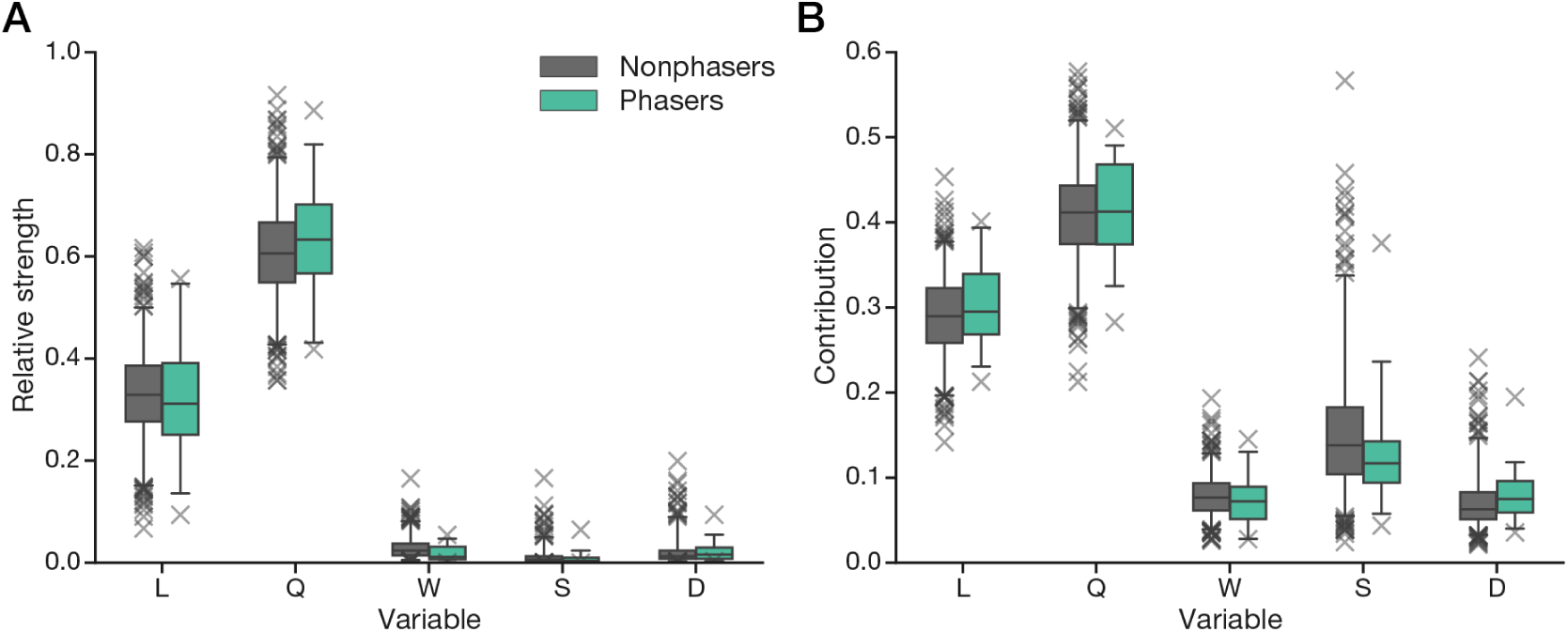
Phaser GLM predictions are driven by spatial predictors. The GLM weights (A) and contributions (B) from the spatial (*LQW*) and trajectory-based (*SD*) variables for nonphasers (*n* = 631 unique cells) and phasers (*n* = 69) are shown in 95% box-and-whisker plots with outliers (×). For phasers, the second-order spatial predictors (*L* and *Q*) are dominant.

Is LQW-SD accurate enough to reproduce spatial activity? The foregoing analysis is predicated on the model’s ability to explain firing patterns. We used LQW-SD spike predictions across the training grid to reconstruct ratemaps (Methods). Quantifying accuracy as the vector cosine similarity between observed and predicted ratemaps, we found phasers (*n* = 69 unique cells; median, 0.994) and nonphasers (*n* = 631; 0.927) to have highly accurate reconstructions (*post hoc* Mann-Whitney *U* = 16,153, *p* = 0.0004). Observed and LQW-SD-predicted ratemaps are shown in Figure 8A-E for the example phasers in Figure 6A-E with overlaid arrows representing the modeled directionality (*β*_*D*_) of each grid section. To verify that LQW-SD also captured strong directional (high DSI) cells accurately, examples of coherent (high DCI) and inhomogeneous (low DCI) directionality are shown in Supplementary Figure 15. Thus LQW-SD provided a high-fidelity account of theta cell firing, including spatial and directional theta cells.

**Figure 8.**
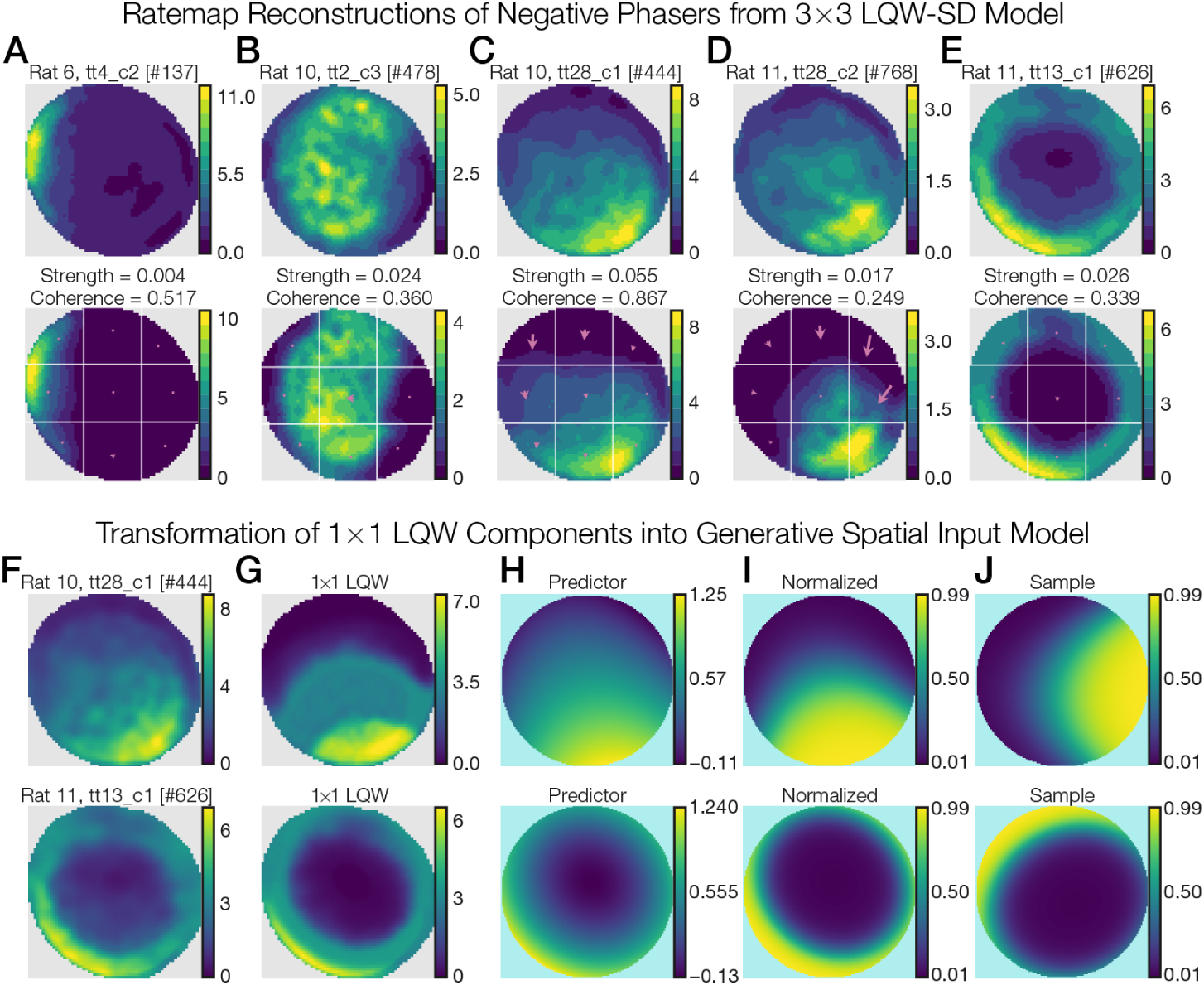
Phaser GLM reconstruction and transformation into generative spatial inputs. (A-E) We show the observed ratemaps (top) and 3 × 3 LQW-SD-predicted ratemaps (bottom) for the phaser examples in Figure 6A-E. Reconstructions were built from spike-count predictions in each grid section (Methods). White lines, model grid boundaries; arrows/dots, normalized GLM directional (*D*) weights; Strength = DSI; Coherence = DCI. (F-J) The steps to form the generative spatial input model are illustrated (Methods). Phasers (F; examples from C and E) are trained in the 1 × 1 LQW model (G), whose linear predictor (H) is normalized to {0, 1} with a sigmoid nonlinearity (I). To generate novel samples (J) from the normalized spatial functions, we added 20% Gaussian noise to the LQW parameters and randomly center-rotated the coordinate frame.

### Competitive phase attractors for flexible spatial synchronization

Can phaser cells synchronize a downstream target to path-integration-like spatial phase codes? We combined our model phasers (equation (3); Table 2; Table 3) with input from a reduced LQW-SD model (equation (1)). The LQW model was trained on the full trajectory (that is, a 1 × 1 grid) without trajectory variables *S* or *D*. The result is a seamless spatial model of biological phaser cell input

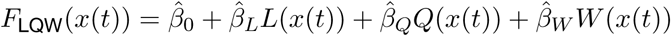

based on trajectory position over time *x*(*t*). LQW training was optimized to wall signals (Supplementary Figure 12) to ensure the minority of boundary-like responses were captured. In our phaser model, spatial functions drive the negative phasers (Figure 2A). To create a generative spatial input model, we selected negative phaser recordings (such as Figure 8F+G) and computed the linear predictors (H), which we normalized to spatial functions on the range {0, 1} (I). We generated novel samples 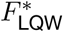 by randomly choosing a spatial function, fuzzing its parameters, and center-rotating the function by a random angle (Figure 8J; Methods). The ramping input (equation (4); Methods) to a model negative phaser thus follows

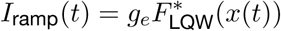

with other parameters unchanged. We simulated 1,000 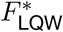 samples driving 1,000 model negative phasers connected (equation (5)) to 1,000 model positive phasers. The simulated phasers (see Supplementary Figures 16+17 for examples) showed a distribution of place-like, gradient-like, and boundary-like responses like the biological phasers but with rate-phase correlations from the model (Figure 1C). Thus model phasers derived from theta cell recordings can help simulate realistic spatial phase synchronization.

**Table 2.**
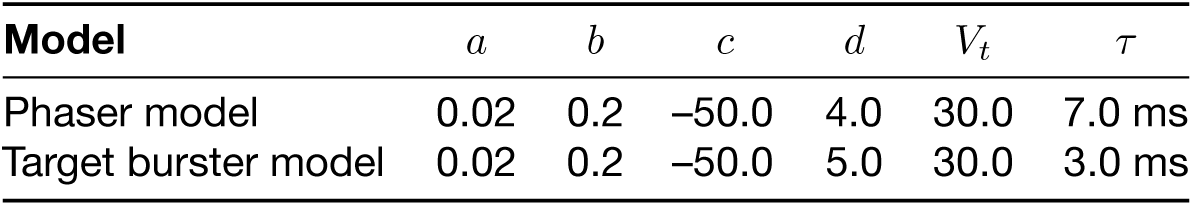
Izhikevich parameters for theta-bursting neuron models.

**Table 3.** Input and conductance parameters for the phaser models.

| Subtype | $g_e$ | $g_\theta$ | $d_{\text{inh}}$ | $E_{\text{inh}}$ | $\tau_{\text{inh}}$ |
| --- | --- | --- | --- | --- | --- |
| Negative | 21.0 | -5.0 | - | - | - |
| Positive | - | 25.0 | 3.0 | -80.0 (mV) | 100 ms |

Do data-driven model phasers support multiplexed spatial synchronization? We first simulated the target burster neuron (equation (6); Table 4) on a 1-hr trajectory (Figure 4A, gray line) without any phaser input. The burst phasemap for this random walk (Figure 9A; peak coherence, 0.486) shows the modulation expected from averaging finite data on a fixed trajectory. We devised spatial phase codes that span both the environment and the theta cycle (Figure 9B) representing path integration along the 45° diagonal. With 2,000 possible phaser inputs, we increased WTA competition to yield 3.5% sparsity (Table 4). The competitively weighted average of joint space-phase distributions (Figure 9C) shows the total phaser input, which reveals a blue band (*π*/2 phase, top), due to positive phasers, alternating with a pink band (*π*, bottom), due to negative phasers. The spatial position of these bands (Figure 9C) tracked the corresponding phase strips in the desired phase codes, indicating that phaser diversity and competitive learning were sufficient to control the input distribution of burst phase across space. Does this phaser input entrain the target burster? With phaser input, the target burster phasemaps (Figure 9D; peak coherence, 0.994, top; 0.973, bottom) reveal highly coherent regions of synchronization to the positive (top) and negative (bottom) phasers that were sharply separated by a narrow band of phase incoherence (darkened area; Supplementary Movie 1). The two synchronization regions were expanded and shifted along the 45° diagonal relative to the input. Both effects are analogous to features of the 1D phase trajectories in Figure 3D+E: The expansion relates to the continuation (horizontally) of burst phase as position moved away from the fixed points; the shift relates to the phasic delay (vertically) between the fixed points and the onset of synchronized target bursting. Can a spectrum of spatial phase codes be learned simultaneously? We simulated 64 target bursters trained on phase codes with varying preferred directions. The same population of model phasers maintained control of the synchronization regions across preferred directions and spatial offsets (Supplementary Movies 2+3). Thus realistic phasers support functionally flexible but dynamically constrained synchronization to spatial phase codes.

**Table 4.** Parameters for the intrinsic theta-bursting neuron used as a synchronization target. WTA values show the percent and number of competitive synapses selected in each model.

| Target model | $I_{\text{const}}$ | $g_{\text{neg}}$ | $g_{\text{pos}}$ | $\sigma$ | WTA |
| --- | --- | --- | --- | --- | --- |
| Target burster (1D) | 12.65 | 1.0 | 2.0 | 0.3 | 20% (50/256) |
| Target burster (2D) | 12.65 | 10.0 | 5.0 | 0.3 | 3.5% (70/2,000) |

## Discussion

We presented network and statistical models that outlined a novel mechanism for anchoring spatial representations in continuous regions of neural synchrony. Simulating the bursting phaser model in 1D, we demonstrated that a simple connectivity motif between theta cells leads to negative and positive rate-phase correlations that can synchronize an artificial step-like phase code to distinct theta phases. We recorded theta cells from hippocampal and subcortical areas in exploring rats and found spatial responses, comprising strong negative (phase precessing) and weaker positive (phase processing) rate-phase correlations, with similar rate and timing dynamics as model phasers. A space-trajectory GLM trained on spike counts showed that trajectory dependence and potential behavioral biases were dominated by pure spatial factors in these cells. Finally, spatial GLM components founded a generative model of environmental inputs to simulate populations of 2D phasers. Sparse competitive weights produced a spectrum of synchronization regions for path-integration-like phase codes across preferred directions and spatial offsets.

**Figure 9.**
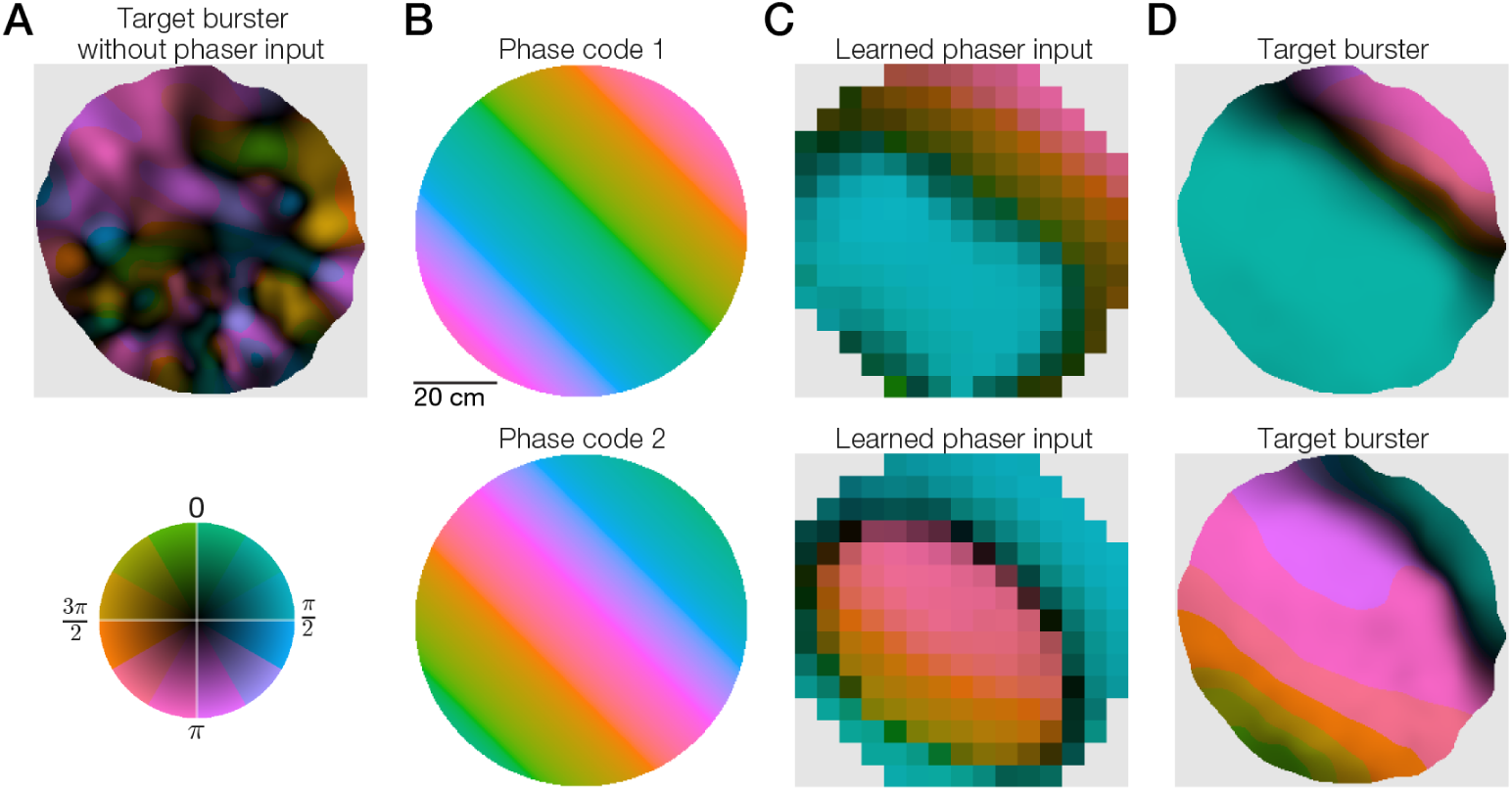
Phaser synchronization of reference points for path integration. We simulated the noisy target burster and 1,000 negative and positive 2D phasers trained on path-integration-like phase codes. (A) Phase-coherence map of burst timing of the target burster without phaser input. Inset, phase-coherence color wheel. (B) For training, two spatial phase codes integrate along the 45° diagonal with different spatial offsets. (C) Phase-coherence maps of competitively-weighted phaser input. (D) Phase-coherence maps of burst timing of the target burster with phaser input. The broad blue/green (top) and pink (bottom) regions represent synchronization by the positive and negative phasers, respectively.

### Relationship to hippocampal place-field theta-phase precession

Phaser timing follows from the notion that stronger input, perhaps representing a sensory value, reduces latency to the next spike. The addition of an oscillating input forms peaks and valleys in time that control when spiking starts (for weak sensory drive above the threshold for silence) and how early the next spike can occur (for strong sensory drive below the threshold for nonstop firing). Mehta *et al* (2002) outlined this mechanism as a conceptual model for how information about position within a place field is encoded in the theta-phase precession of place cell firing (O’Keefe & Recce, 1993). In that theory, the sensory input comprises synaptic drive from the place cell network. That network input forms an asymmetric ramp-like input after learning and exploration (Mehta et al., 1997, 2000), allowing place cells to monotonically precess across any traversal of the place field (Schmidt et al., 2009). The additional plasticity, neuronal, and network effects supporting place-field phase precession may contribute to diverse functions including place field formation, spatial precision, and sequence learning for navigation and memory (O’Keefe & Recce, 1993; Skaggs et al., 1996; Jensen & Lisman, 1996; Wallenstein & Hasselmo, 1997; Levy et al., 2005; Dragoi & Buzsáki, 2006; Feng et al., 2015). The phaser model, however, is driven by a layer of sensory inputs represented as functions of space, without learning or network inputs. The phaser precesses/processes as the spatial input increases/decreases, creating an experience-independent temporal code that maps phase directly to an isocontour of its input function. This relative mechanistic and functional simplicity makes phasers parametrically robust and potentially prevalent in brain areas with oscillations and spatial/sensory inputs. This robustness allowed our models to be broadly tuned with few parameters (Table 2; Table 3; Methods). Thus a simple mechanism may enable failsafe implementations of an important neurocomputation.

### Competitive burst synchronization as spatial phase reset

Conversion between rate and phase codes may be an important neurocomputation for spatial/sensory feedback. The Mehta mechanism can yield highly correlated rate and phase, which raised the question whether temporal codes carry information distinct from firing rate. In place-field phase precession, there is evidence that phase and rate distinctly encode distance and speed (Huxter et al., 2003), which inspired the oscillatory interference theory whereby path integration is computed with a phase code (O’Keefe & Burgess, 2005; N. Burgess, 2008; Hasselmo, 2008). This theory described neural oscillators (VCOs) that integrate speed-modulated direction inputs with changes in phase, allowing conversion via neural synchrony into the firing-rate representations of grid and place cells (Blair et al., 2008, 2014). VCOs were originally conceived as dendritic oscillations, which allowed for simple reset mechanisms (N. Burgess, 2008; Hasselmo, 2008) but disallowed the necessary independence of multiple oscillators (Remme et al., 2010). While some theta cells in rats demonstrate the directional tuning of burst frequency necessary for VCOs (Welday et al., 2011), other species such as fruit bats have grid cells without continuous theta oscillations (Yartsev et al., 2011). Additionally, the critical theoretical problem for VCO codes is accumulating errors in self-motion inputs, including head-direction inputs that may not align with the animal’s movement direction, and noise from biophysical variance in the oscillators. These errors lead to random drift of the spatial code within its reference frame or teleportation of the code to a different environment.

Proposed stabilizing mechanisms include network synchronization, ring attractors, oscillator redundancy, and/or coupling with continuous attractors in the grid cell network (Zilli & Hasselmo, 2010; Hasselmo & Brandon, 2012; Bush & Burgess, 2014; C. P. Burgess & Burgess, 2014; Blair et al., 2014). Those models did not examine intrinsic neuronal bursting, as we do here, but it may also help stabilize phase-coding neurons by regulating burst frequency via resonance (Lisman, 1997; Izhikevich, 2007) and reducing the impact of intra-burst spike-count or spike-interval variance on the phase code.

Stabilization alone does not address how environmental cues reset or calibrate the phase code within an absolute spatial reference frame. Simple phase models of hippocampal remapping demonstrated that regular calibration from sensory cues eliminates phase drift over long timescales (Monaco et al., 2011). The phaser mechanism that we study here is a rate-to-phase conversion that may provide the synchronous feedback signal needed for that calibration. Phasers take direct spatial inputs that modulate rate and phase together such that rate controls feedback gain (to downstream targets) and burst phase indicates the input-mapped phase target. This mechanism does not require continuous theta oscillations: Transient bouts of theta would propagate brief synchronizing bursts. The intrinsic bursting dynamics (equation (3); Izhikevich, 2007) reproduced some characteristics of parahippocampal theta cells such as skipping or alternating theta cycles (Deshmukh et al., 2010; Brandon et al., 2013). With competitive weights, phasers were able to collectively synchronize a noisy theta-bursting neuron to both 1D (Figure 3) and 2D (Figure 9) spatial phase codes. These simulations demonstrated that weak clock signals (that is, phasers with intra-burst spike variance and cycle skipping) can collectively counteract accumulated phase errors in a neuronal oscillator (cf. Rossant et al., 2011). The role of bursts in facilitating interareal transmission and synaptic plasticity (Lisman, 1997; Csicsvari et al., 1998) suggests bursting could be critical to an experience-dependent feedback loop. Thus the calibration of the spatial metric may depend on burst phase-synchronization intermediating rate-based representations.

### A conversation between calibration and integration

Our phaser simulations present two hypothetical populations (negative, positive) with a minimal connectivity scheme (Figure 2A). This modeling choice restricted the granularity of phase fixed-points that could be synchronized: More complex connectivity patterns may increase the diversity of preferred phases available for spatial synchronization. In the model, negative/positive phasers demonstrated a pattern of strong/weak rate-phase correlations and trough/peak theta preference (Figure 1C). Extracellular recording precluded subthreshold evidence for the mechanism, but the dynamics of identified biological phaser cells corroborated this model phaser pattern (Figure 5). The GLM model of biological phaser firing further corroborated phasers’ nondirectionality or directional isotropy, which is a general requirement for stable path integration systems (Issa & Zhang, 2012). Thus biological phasers might support broadly, not finely, tuned spatial phase attractors (cf. Supplementary Movie 1).

How can broadly-tuned calibration support finely-tuned integration? One possibility is that calibration is selectively activated after learning. Learning requires synchrony with phasers, but the burst frequency of VCOs increases with movement along the preferred direction (N. Burgess, 2008; Welday et al., 2011). The subset of VCOs with preferred directions orthogonal to the animal’s movement direction will phase synchronize with the shared theta rhythm and, correspondingly, the phaser population. Thus the orthogonal subset will optimize competitive learning for phasers over repeated theta cycles in the same location; this subset evolves as the animal explores the environment. Calibration in the familiar environment likewise requires synchrony with phasers, but it must be interdigitated with path integration (Monaco et al., 2011) perhaps mediated by discrete attentive behaviors during pauses in locomotion such as head scanning (Monaco et al., 2014). Without this interplay, the spatial precision of the phase code would be bounded by the broad tuning of the phase attractor. Ring attractor organization of VCOs (Blair et al., 2014; C. P. Burgess & Burgess, 2014) could have particular benefits during initial learning and online calibration: Respectively, to ensure the existence of an orthogonal subset for any movement direction, and to propagate the calibrated phase throughout the path integrator network via intrinsic connectivity. Selective switching between calibration and integration could be driven by accumulating error or mismatch signals, possibly mediated by grid cells (Blair et al., 2014; Rennó-Costa & Tort, 2017). Further studies should characterize how this phase code “conversation” might support the brain’s spatial metric.

### Hippocampo-septo-entorhinal feedback loop for the spatial metric

Biological phasers were found in hippocampus and lateral septum, but not thalamic sites. Lateral septum, with the bulk of the recording data (Table 1), is interconnected with hippocampal CA1/CA3 and pacemaker networks of the medial septum (Jakab & Leranth, 1995). These cells are well-placed to combine theta oscillations and spatial inputs (Takamura et al., 2006), as required for the phaser mechanism, and to participate in subcortical theta-rhythmic feedback and regulatory circuits (Leranth et al., 1999; Luo et al., 2011; Sartor & Aston-Jones, 2012; Ruan et al., 2017). The spatial representations of biological phasers included small place-like fields, broad linear gradients, and border/border-place responses. This spatial variance could be driven by diverse inputs and connectivity patterns including single or multiple place cells, object-place cells (Deshmukh & Knierim, 2011; Tsao et al., 2013), or border cells (Lever et al., 2009; Savelli et al., 2008; Solstad et al., 2008). While our data-driven simulations of 2D phasers showed constrained spatial tuning (as discussed above), training with VCO-like phase codes flexibly produced border-aligned regions of phase synchronization across preferred directions (for example, Figure 9D, top, pink). If path integration is calibrated by a phaser mechanism, then these learned regions could contribute to the role of border visits in correcting (or distorting) the spatial metric carried by grid cells (Hardcastle et al., 2015; Stensola et al., 2015). Thus robust temporal neurocomputations may help anchor neural spatial maps to the outside world.

## Materials and methods

### Bursting models

We define a quadratic integrate-and-fire model (Izhikevich, 2003) of intrinsic bursting with a fast variable for the spiking limit cycle (*V*) and a slow adaptive variable for terminating bursts (*u*). The dynamics follow

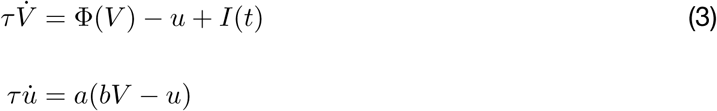

where *I*(*t*) is a cell-specific time-varying input, Φ(*V*) = 0.04*V*^2^ + 5*V* + 140 is a quadratic nonlinearity for spike initiation, *a* and *b* control adaptive feedback, and *τ* sets a shared time-scale for spiking and bursting (in addition to the time constants implicit in Φ(*V*) and *a*). Whenever *V* > *V*_*t*_, a spike is recorded, *V* is reset to *c*, and *u* is incremented by *d*. Bursting parameters are listed in Table 2. While *V* is approximately millivolt scale, we treat this system as a qualitative, not biophysical, model for which the parameters are in arbitrary units.

For negative phasers, we set the time-varying input (equation (3)) to the combination

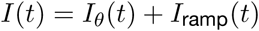

of sinusoidal theta inhibition (for inhibitory gain *g*_*θ*_ < 0)

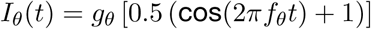

and direct ramping excitation (for excitatory gain *g*_*e*_)

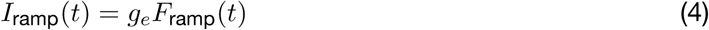

where the ramping input function *F*_ramp_(*t*) has range {0, 1}.

The positive phasers have theta gain *g*_*θ*_ > 0 and follow equation (3) with negative-phaser input

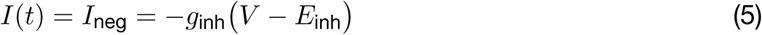

where *g*_inh_ is a slow inhibitory conductance

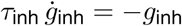

that is incremented by *d*_inh_ with every pre-synaptic spike (Table 3).

The target bursters have a shorter time-constant (*↓τ*) and lower burst excitability (*↑d*; Table 2). In place of equation (3), the fast variable follows

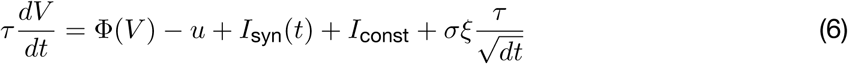

where normalized white noise *ξ* is controlled by gain *σ*, and *I*_syn_(*t*) is total synaptic drive from the phaser network

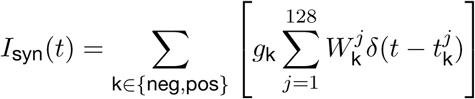

where *g*_neg_/*g*_pos_ are subtype-specific feedback gains (Table 4), *W*_neg_/*W*_pos_ are the phaser weight vectors (for example, Figure 3B), and *t*_neg_/*t*_pos_ are most-recent-spike vectors. Constant input current was tuned (*I*_const_, Table 4) so that the intrinsic burst rate, without noise or synaptic input, was close to reference theta frequency (7.519 *s*^−1^ compared to *f*_*θ*_ = 7.5 *s*^−1^).

### Spiking simulations

Spiking neuron and network models were implemented in the equation-based Brian simulator (Goodman & Brette, 2008). Simulations were integrated in 1-ms timesteps. Phaser layers and the target burster without noise were evolved with Runge-Kutta 4th-order integration; the target burster with noise used the forward Euler method. Burst timing in simulations was determined as spike times following interspike intervals ≥25 ms.

For 1D spatial simulations, local tuning functions were Gaussian functions with bandwidth 1/64 normalized to {0, 1} and centered at 64 evenly-spaced positions from 0 to 1. Each long-range tuning function was 1 minus a local tuning function. The gain of phaser input onto the target burster (Table 4) was manually tuned for visually matched ‘middle of the road’ synchronization at both fixed points.

For 2D spatial simulations, phase-code target gratings had spatial period 80 cm so that one cycle covered the environment. Phaser gain onto the target burster (Table 4) was manually tuned to roughly equalize the size of negative and positive synchronization modes across different reference phases.

### Competitive learning

Based on 1-hr training simulations, we generated joint space-phase distributions from phaser spikes: 15 × 36 (*x*, *ϕ*) bins for 1D simulations; 15 × 15 × 36 (*x*, *y*, *ϕ*) bins for 2D simulations. The phase-code target was either directly specified as a binary array for 1D simulations or binned from a spatial grating function for 2D simulations. We computed the vector cosine similarity between the phaser distributions and the target as the basis for the phaser synaptic weights. To determine competitive weights, we chose the WTA% negative and WTA% positive phasers (Table 4) with the highest similarities and normalized those similarities to {0, 1} via [(similarity − min) / (max − min)]. Unselected weights were set to 0. Weighted-average phaser inputs (Figure 3C; Figure 9C) were computed as the product-sum of the weight vector with an array of all space-phase distributions.

### Subjects and surgery

Male Long-Evans rats (350–400 g) were individually housed and kept at 85% of *ad libitum* weight. They were trained over 5 d to forage for food pellets in an enclosed environment. Under deep isoflurane anesthesia, rats were chronically implanted with tetrode arrays targeting (across rats) the septum, dorsal hippocampus, anterior thalamus, midbrain, and/or other subcortical areas. Each rat was implanted with 16 tetrodes (64 electrode channels) that were grouped into four independently drivable bundles of four tetrodes each. All experiments were conducted in accordance with the U.S. National Institute of Health Guide for the Care and Use of Laboratory Animals (NIH Publications No. 80–23), and were approved in advance by the animal subjects review committee at the University of California, Los Angeles.

### Theta cell recordings

Data collection methods including conduct of recording sessions, video tracking analysis, and single-unit acquisition have been described previously (Welday et al., 2011). The phase of the septohippocampal theta oscillation was quantified from the LFP signal on a reference electrode in the hippocampal fissure. In one subject (rat 11), a strong theta cell was used as phase reference instead of the LFP signal and was not otherwise included in data analysis. All data for analysis was filtered for linear movement speeds >5 cm/s.

### Adaptive spatial maps

To handle large variance in spatial data density from long recordings, we computed spatial maps with adaptive scaling kernels. We used a KD-tree algorithm to generate a nearest-neighbor model of the data points for the map. For every pixel to evaluate, we found the enclosing radius of the nearest 4% of data points. If the radius was <8% or >30% of the arena diameter, then it was fixed at 8% or 30%, respectively. A Gaussian kernel set weights for each data point in this evaluation radius. For ratemaps, we computed weighted averages of trajectory data and spike data to create occupancy and spike density maps; dividing the spike density by the occupancy map produced the ratemap. For phasemaps, we computed weighted mean resultant vectors from which we retrieved the phase average and variance; the phase average was used for phase-only maps and the variance was normalized into a coherence mask for the phase-coherence maps.

### Theta-rhythmic analysis

The rhythmicity index and burst-frequency estimates were derived from spike-timing autocorrelations. We adaptively smoothed 128-bin 0.5-s correlograms to find stable estimates of the first trough and first (non-central) peak of the correlograms. Rhythmicity was calculated as the ratio [(peak − trough) / peak]. Burst-frequency was calculated as the average of the first-peak mode estimate and an estimate based on a weighted-average of the first-to-second-trough correlations.

The theta modulation index was computed from a 10° binned phase histogram on {*−π, π*}. We circularly convolved the histogram with a 10° bandwidth Gaussian kernel for smoothing. Theta modulation was calculated as the ratio [(max − min) / max] of the smoothed histogram.

### Rate-phase regressions

We implemented the method of Kempter *et al* (2012) for computing circular-linear regressions with stable estimates of the correlation coefficient and *p*-value. Unreported *p*-values were arbitrarily close to 0. This method was used for all rate-phase regression lines and correlation values. To compute total phase shift, we multiplied the rate-phase regression slope by the width of the range of firing rates in the ratemap.

### Information-theoretic measures

We computed spatial phase information *I*_phase_ as the mutual information between phase (*ϕ*) and position (*x*)

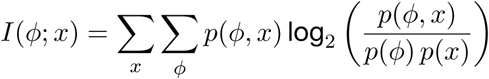

based on joint space-phase distributions of spikes binned into 15 × 15 × 36 (*x*, *y*, *ϕ*) arrays. This measure yields information in units of bits. We permuted spike phases 1,000 times to calculate *p*-values.

We computed spike information measures based on Skaggs’ (1993) formulation

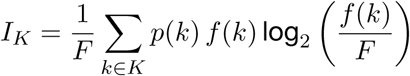

where *K* is position, direction, or speed of the trajectory; *p* is the occupancy density; *f* is a firing-rate function; and *F* is the mean firing rate. Position was binned into 15 × 15 arrays on {0, 80} cm along the *x* and *y* axes; direction into 36 bins on {0, 2*π*}; and speed into 18 bins on {5, 50} cm/s excluding bins with <3 s occupancy. These measures yield information rates in units of bits/spike. We randomly shifted-and-wrapped spike trains with 20 s minimum offsets and reinterpolated trajectory data 1,000 times to calculate *p*-values.

### Movement modulation

The direction modulation index was computed as the ratio [(max − min) / max] of a smoothed firing-rate function of movement direction. Average firing rates in 36 direction bins on {0, 2*π*} were circularly convolved with a 10° bandwidth Gaussian kernel. The speed modulation index was computed as the ratio [(max − min) / max] of a firing-rate function of speed. Average firing rates were calculated for 14 bins on {5, 40} cm/s excluding bins with <8 s occupancy.

### GLM training

Ridge regression models were trained on 9 scalar predictors representing the vector components of the 5 model variables: *L* = (*x*, *y*), *Q* = (*x*^2^, *y*^2^, *xy*), *W* (scalar), *S* (scalar), and *D* = (*u*_*x*_, *u*_*y*_). The wall predictor *W* was a sigmoid proximity signal [1/(1 + exp(−*k*(*r − w*_0_)))] for radius *r* from arena center, *k* = 0.5, and *w*_0_ = 30 cm. *S* was linear trajectory speed. *D* was the unit vector along the movement direction. Training samples were 300-ms bins and predictors were interpolated at the midpoint of each bin. Each predictor was standardized by subtracting its sample mean and dividing by its sample standard deviation. The response variable was the log spike-count *Y* for each bin, which makes this a Poisson-distributed GLM. The trajectory was divided into equal-sized 2 × 2 or 3 × 3 grids based on data limits. For each grid section, the GLM was trained on all samples inside the section based on the interpolated (*x*, *y*) position. Estimated model intercepts and coefficients for each recording and grid section were stored for analysis (or for the reduced LQW generative model). To regularize the model, tuning parameter *α* determined the *ℓ*_2_-norm penalty for least-squares optimization

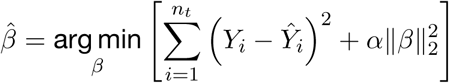

where *n*_*t*_ was the number of training samples. We maximized model directionality (or, similarly, the boundary response *W* for the LQW generative model) by choosing

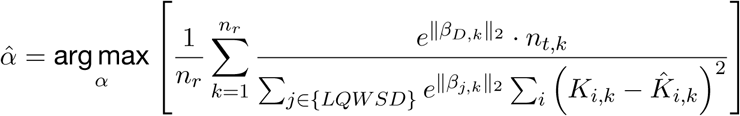

which maximizes (over *n*_*r*_ = 1,073 theta-cell recordings) the softmax directional coefficients while minimizing mean squared error (MSE) of spike-count (*K* = exp(*Y*)) predictions (Supplementary Figure 12). The value *α* = 1.2496 from the 2 × 2 model was used for analysis because of higher likelihood, lower MSE, lower penalty, and complete wall contact across grid sections compared to the 3 × 3 model.

### GLM analysis

The relative strengths of GLM variables were computed as normalized vector norms

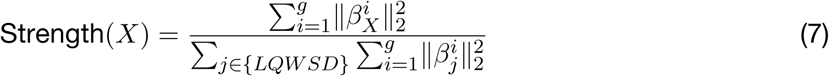

for variable *X* ∈ {*L, Q, W, S, D*} across *g* grid sections. Thus DSI was computed as Strength(*D*) and DCI was computed as 1 minus the normalized circular standard deviation of the *β*_*D*_ vector angles across the grid. We computed variable contributions similarly to equation (7) but with maximum linear predictors (equation (2)) instead of coefficient vector norms. The sum across variables for both relative strength and contribution was normalized within recordings and then averaged by unique cell (Figure 7). Grid matrix plots (Supplementary Figure 14A+C) show these values without the grid summations (equation (7)).

To reconstruct ratemaps, we used the midpoints of grid-specific training samples to predict spike counts from the model for each grid section. We collated the counts and sample positions across grid sections to reconstitute a complete dataset for generating the ratemap.

To create the LQW generative model, we used a COBYLA search to find the arena-bounded minimum and maximum of the linear predictor for each recording. We normalized the LQW parameters to {0, 1} and applied a clipping sigmoid [1/(1 + exp(−10 (*f −* 0.5)))] to smoothly enforce the range of the resulting spatial function. To sample the generative model, we randomly selected a negative phaser’s spatial function, added 20% Gaussian noise to its LQW parameters, and rotated the function about the center by a random angle.

### Software

Modeling and analysis was performed using a custom python package that depends on the open source ecosystem: numpy, scipy, matplotlib, seaborn, pandas, scikit-learn, pytables, Brian2, and other libraries. The source code and a complete specification of the python environment is available at https://doi.org/10.6084/m9.figshare.5552467.

## Acknowledgments

This work was supported by the CRCNS grant NIH R01MH079511 to K.Z., H.T.B., and J. J. Knierim. A Johns Hopkins University Science of Learning Institute award supported J.D.M. during the writing of the paper.

### Author Contributions

J.D.M. analyzed data, developed models, performed simulations, and wrote the paper. H.T.B. conducted experiments and collected data. H.T.B. and K.Z. conceived the idea for the project and provided scientific guidance. K.Z. provided mathematical guidance. All authors read and commented on revisions of the manuscript.

---

**Supplementary Figure 1.**
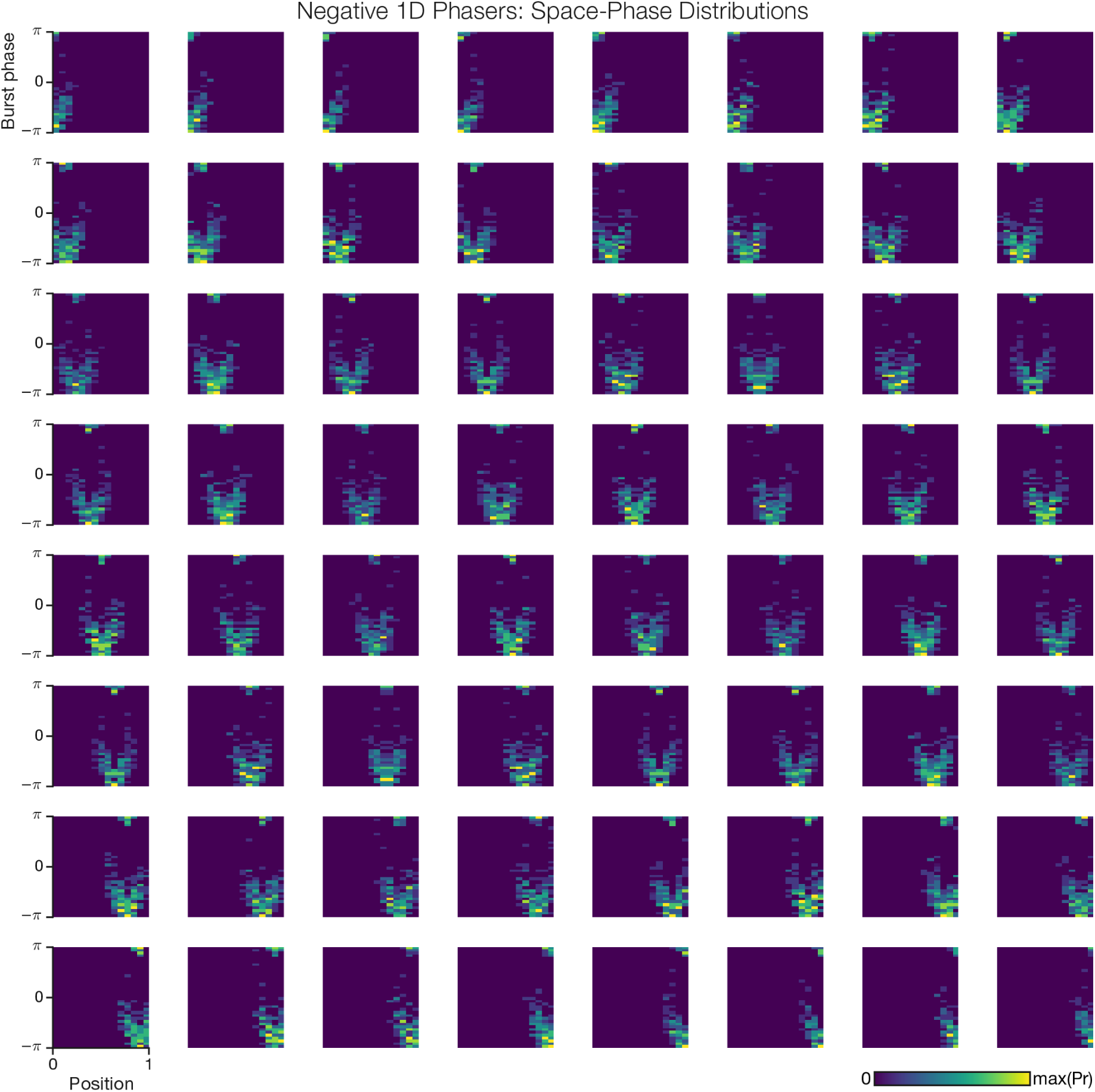
Spatiotemporal activity of 1D spatial network: Negative phasers. This grid shows, from left-to-right and top-to-bottom, the joint space-phase distributions for all negative phaser cells driven by a local tuning function in the network (Figure 2A) from *x* = 0 to *x* = 1. The middle plot corresponds to the top left of Figure 2B.

**Supplementary Figure 2.**
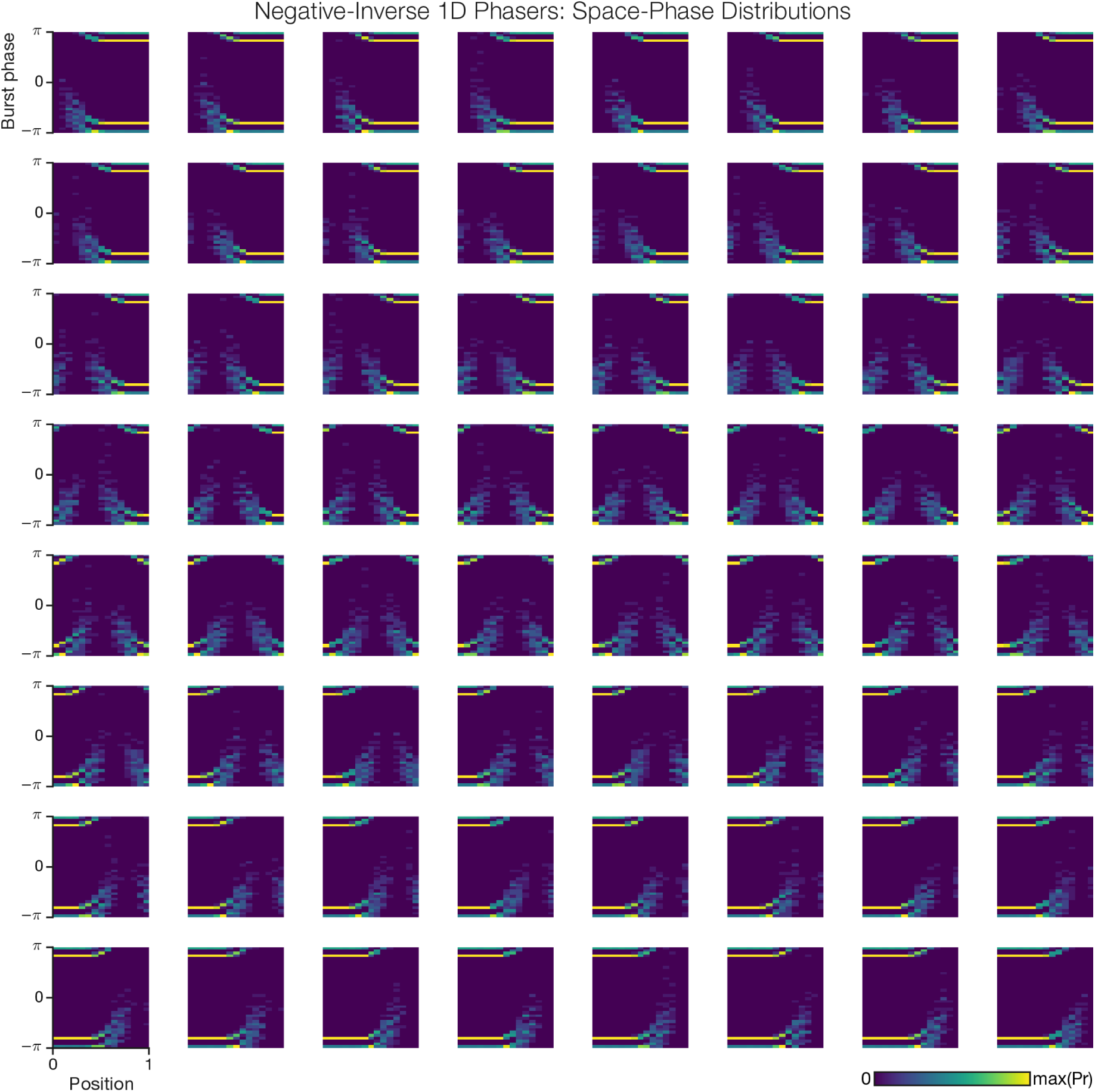
Spatiotemporal activity of 1D spatial network: Negative/Inverse phasers. This grid shows, from left-to-right and top-to-bottom, the joint space-phase distributions for all negative phaser cells driven by an inverse (long-range) tuning function in the network (Figure 2A) from *x* = 0 to *x* = 1. The middle plot corresponds to the bottom left of Figure 2B.

**Supplementary Figure 3.**
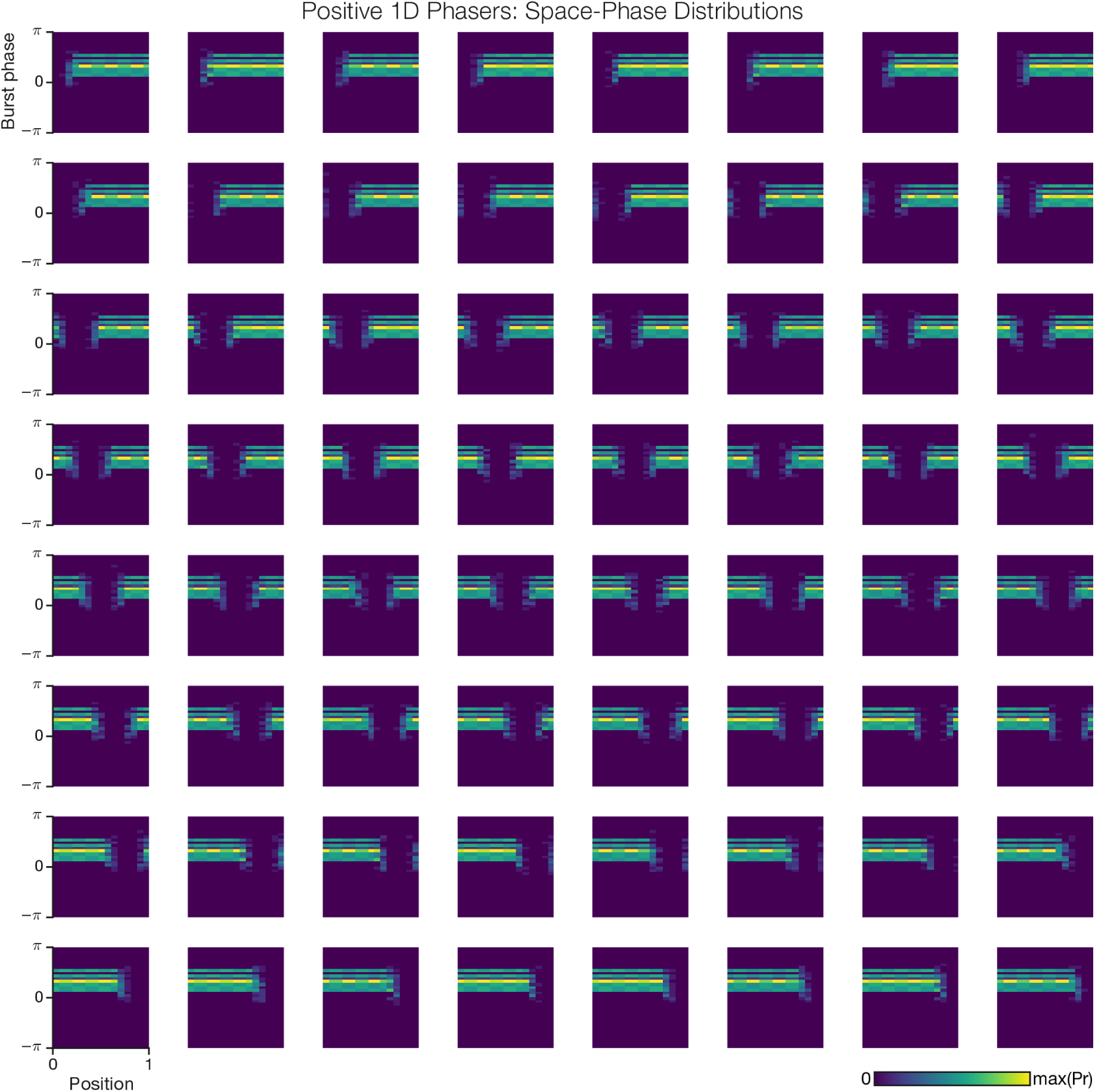
Spatiotemporal activity of 1D spatial network: Positive phasers. This grid shows, from left-to-right and top-to-bottom, the joint space-phase distributions for all positive phaser cells suppressed by a local negative phaser in the network (Figure 2A) from *x* = 0 to *x* = 1. The middle plot corresponds to the top right of Figure 2B.

**Supplementary Figure 4.**
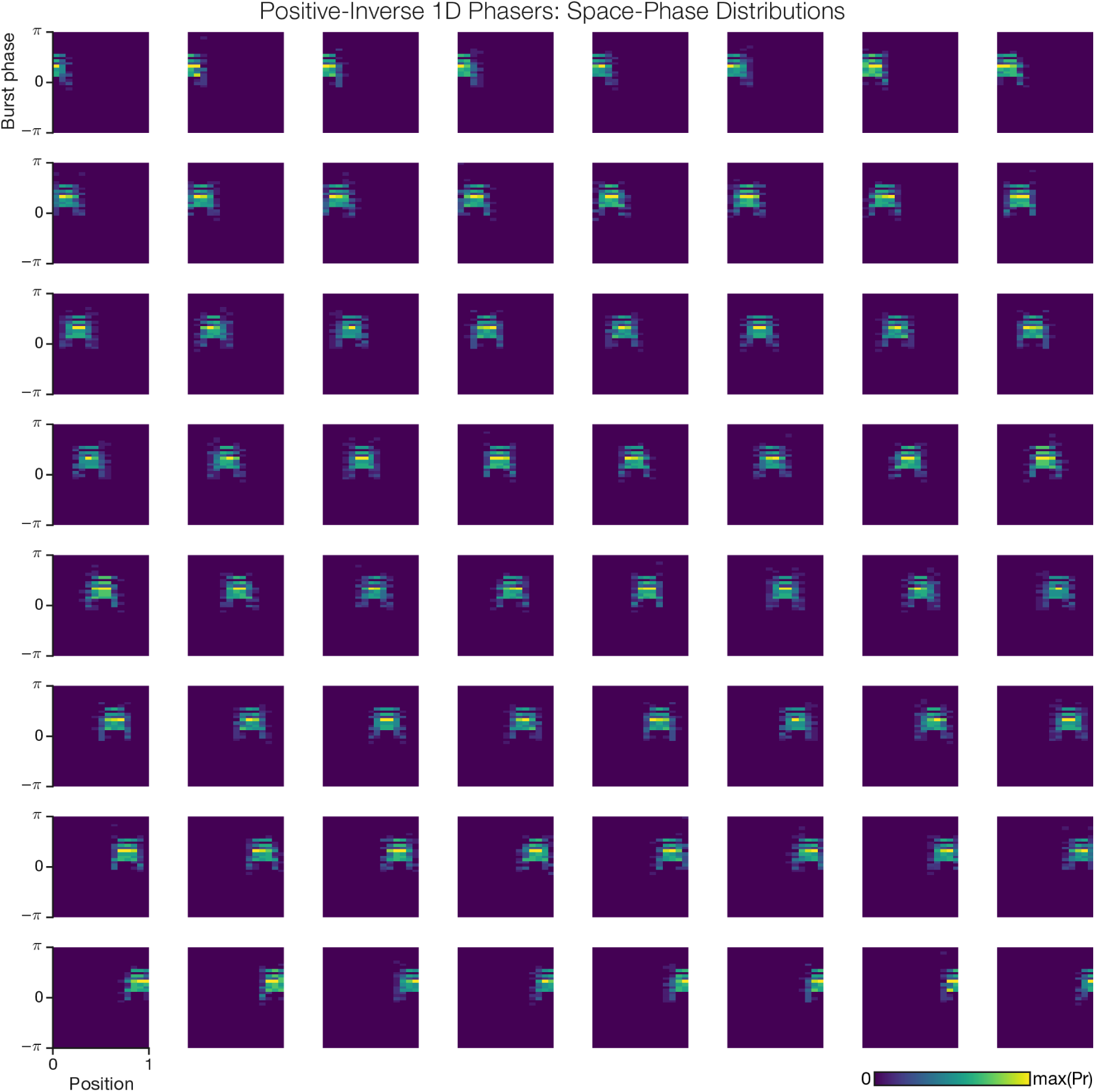
Spatiotemporal activity of 1D spatial network: Positive/Inverse phasers. This grid shows, from left-to-right and top-to-bottom, the joint space-phase distributions for all positive phaser cells suppressed by an inverse (long-range) negative phaser in the network (Figure 2A) from *x* = 0 to *x* = 1. The middle plot corresponds to the bottom right of Figure 2B.

**Supplementary Figure 5.**
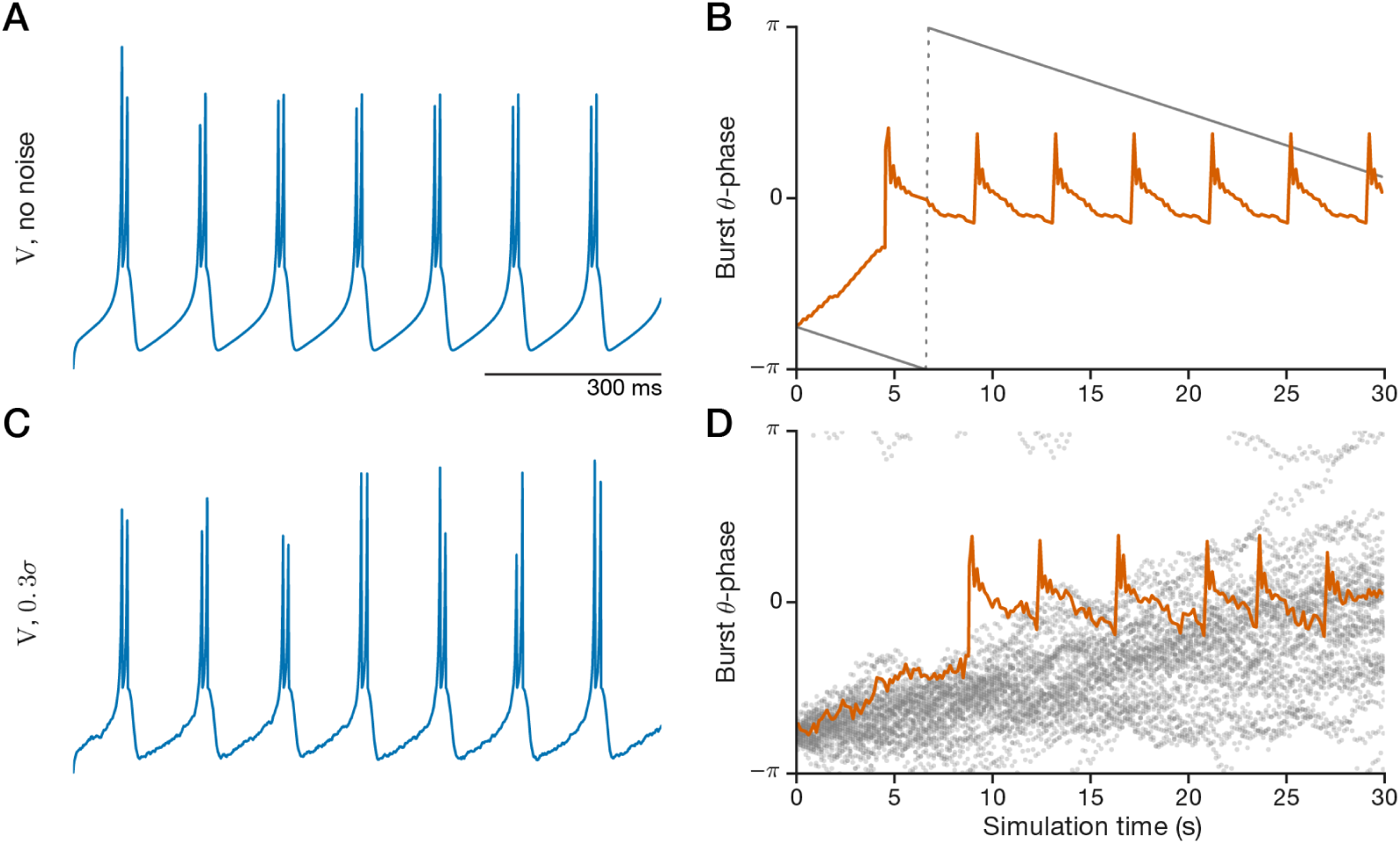
Theta-bursting neuron with phase noise: drift and pulse synchronization. An intrinsic bursting model (equation (6); Izhikevich, 2007) was tuned with constant input (Table 4) to fire doublet bursts (A) close to the reference theta frequency, 7.5 Hz. The difference between the reference and the actual burst rate, 7.519 bursts/s, means that the cell’s theta phase (B) slowly drifts (precesses) over time (gray line). To test whether this cell can be phase-synchronized by periodic stimulation, we simulated an instantaneous pulse (*V* ← *V* + 15mV) every other theta cycle at zero phase. The pulse-synchronized theta cell (B, orange line) monotonically processes toward zero phase and then (around *t* = 5 s) discontinuously jumps past zero phase before slowly precessing to just before zero phase. This dynamic, of jumping forward and precessing back, repeats (around *t* = 9 s) and continues stereotypically. This sawtooth pattern encapsulates the theta-synchronization dynamics of this cell. For our phaser synchronization simulations, we added phase noise to this ‘target burster’ cell (equation (6)) to show that the synchronization can overcome intrinsic noise. We chose a noise level (Table 4) that preserved the cell’s theta bursting (C, same as Figure 3A, inset) but largely randomized its burst theta-phase over the 30-s simulation (D, gray circles, 36 trials). With phase noise, the pulse stimulation reproduced the sawtooth pattern of synchronization (D, orange line).

**Supplementary Figure 6.**
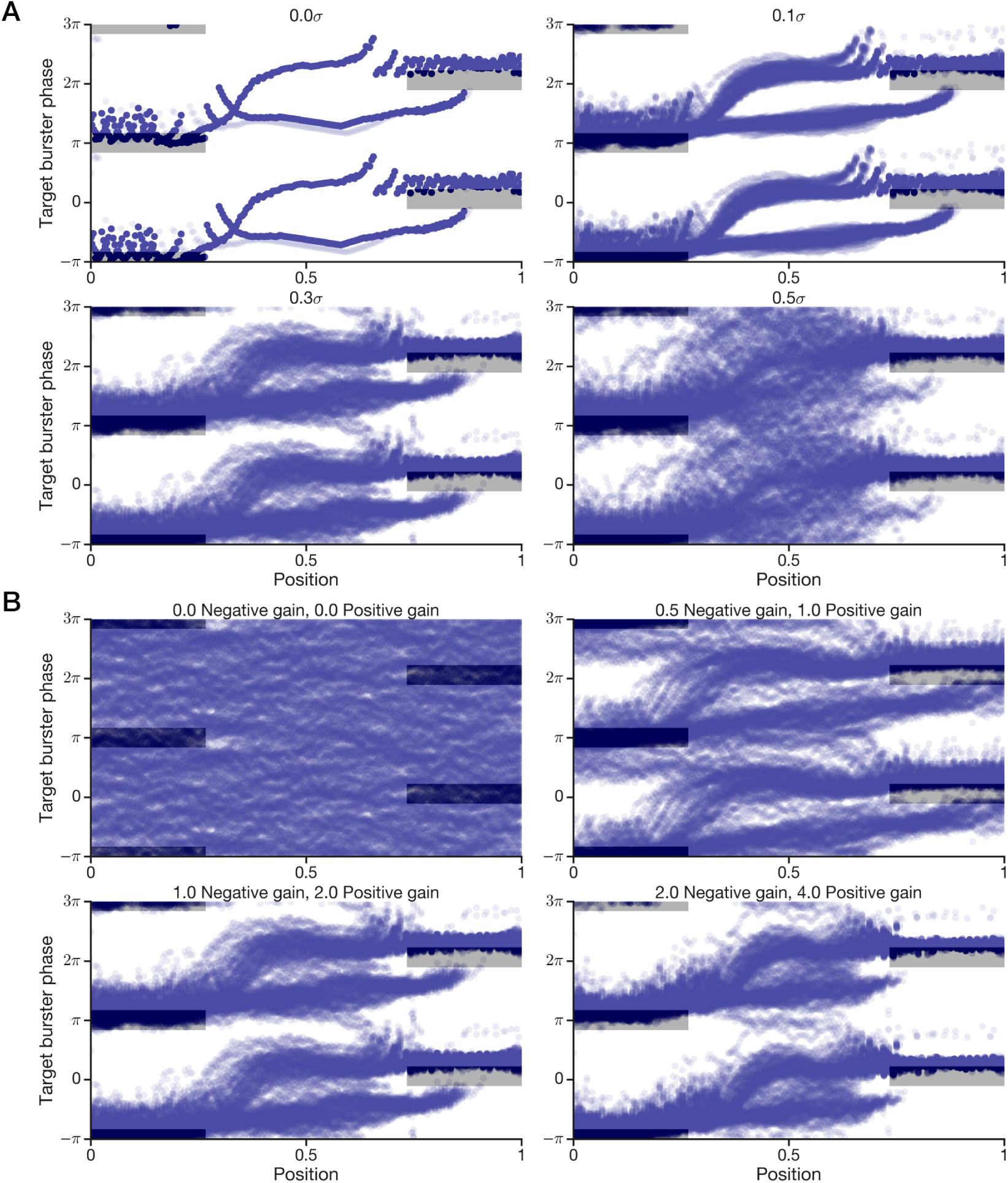
1D phaser synchronization across noise and gain levels. Here we show additional 1-hr simulations of the 1D spatial synchronization network shown in Figure 3. (A) With the gain from the phasers fixed (Table 4), simulations with 0.0*σ*, 0.1*σ*, 0.3*σ*, and 0.5*σ* noise levels demonstrate that the phase-code fixed points remain functional at various noise levels. (B) With the noise level fixed at 0.3*σ*, the effect of zero phaser gain (top left) can be compared to weaker (top right) and stronger (bottom right) levels of phaser gain. Weak phaser gain (top right) still synchronizes the target burster, but the phase trajectories are extended due to the slower development of phase locking on approaches toward *x* = 0 and *x* = 1.

**Supplementary Figure 7.**
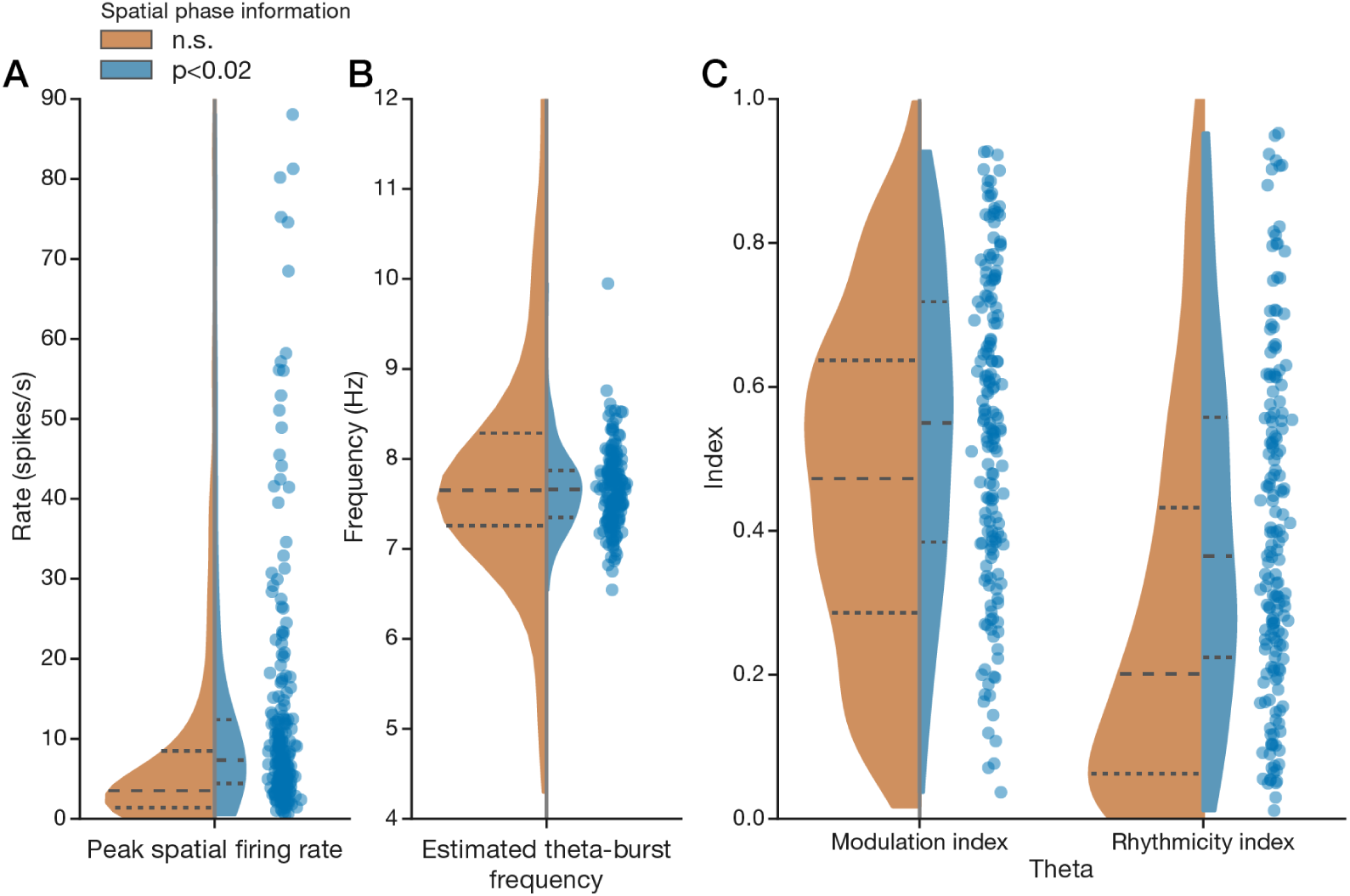
Spatial theta cells are rhythmically normal. For comparison, we show distributions for theta cell recordings split into groups with non-significant (n.s.; ‘nonspatial’; *n* = 840) or significant (*p* < 0.02; ‘spatial’; *n* = 233) spatial phase information (Methods). Gaussian kernel-density estimates (using Scott’s bandwidth rule) of splits are normalized by group size and show medians (long-dash lines) and quartiles (short-dash lines). Scatter points are additionally shown for the spatial data. Peak firing rate (A) and autocorrelogram-based estimates of theta-burst frequency (B; Methods) are similarly distributed for nonspatial and spatial recordings. Theta indexes for modulation and rhythmicity (C; Methods) show that spatial recordings are distributed slightly higher, but this is likely due to the lower modes evident for non-spatial recordings which may consist of borderline or non-theta cells. Thus spatial theta cells have similar firing rate and oscillatory characteristics to theta cells in our dataset.

**Supplementary Figure 8.**
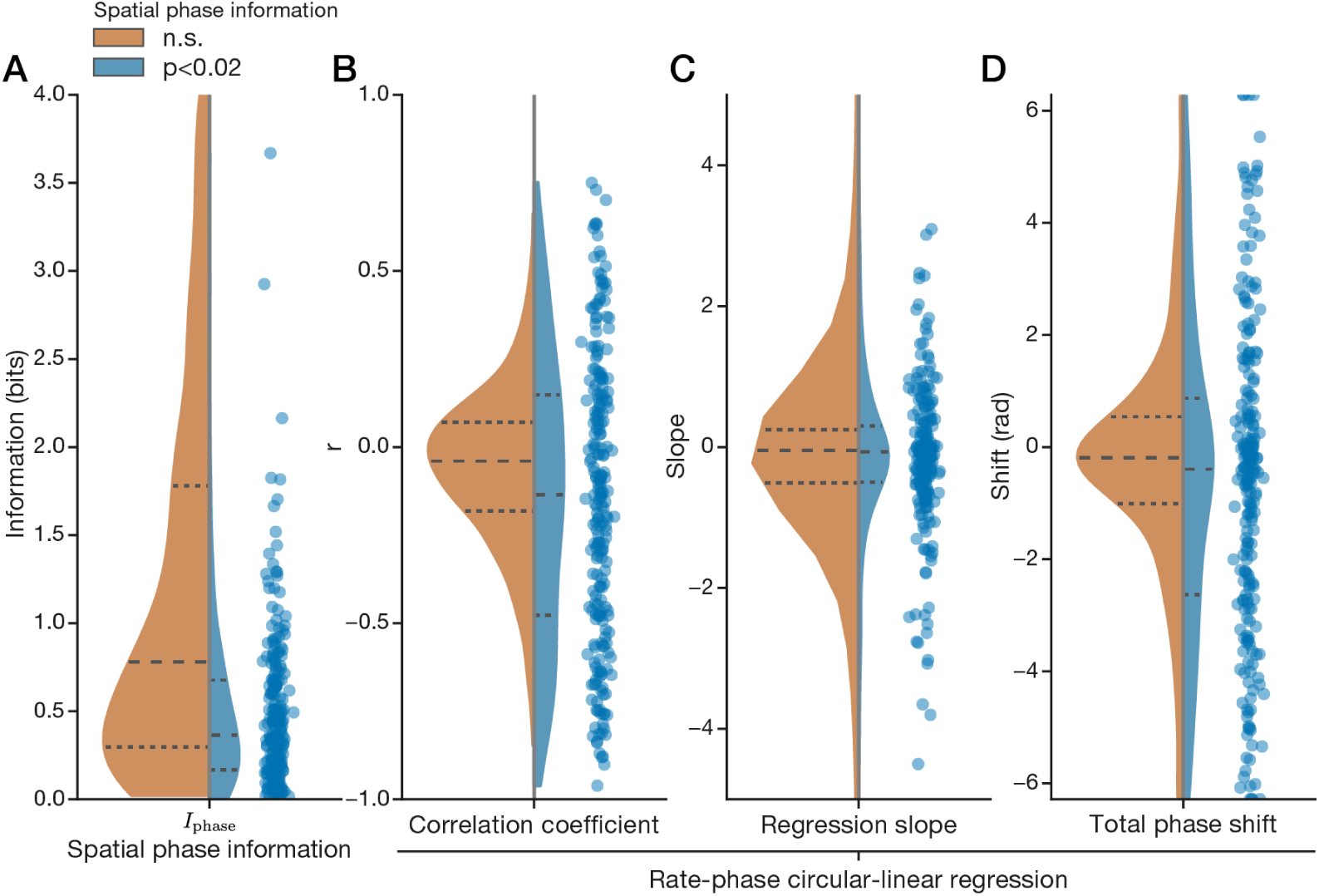
Spatial theta cells have broadly distributed rate-phase correlations. Similar to Supplementary Figure 7, we show distributions of theta cell recordings split into groups with non-significant (n.s.; ‘nonspatial’; *n* = 840) or significant (*p* < 0.02; ‘spatial’; *n* = 233) spatial phase information (Methods). (A) *I*_phase_ for spatial cells has median 0.36 bits/spike (long-dash line) with a positively skewed distribution and wide range. (B-D) Circular-linear regressions of average phase onto average rate based on spatial map pixels. Nonspatial recordings were distributed around zero. Estimates for correlation coefficient (B) and total phase shift (D; Methods) show broader distributions for spatial than nonspatial cells: Compare quartiles (short-dash lines) and fatter tails reflecting excess negative/positive correlations. Total phase shift (D) is computed as a rate-normalized slope (C).

**Supplementary Figure 9.**
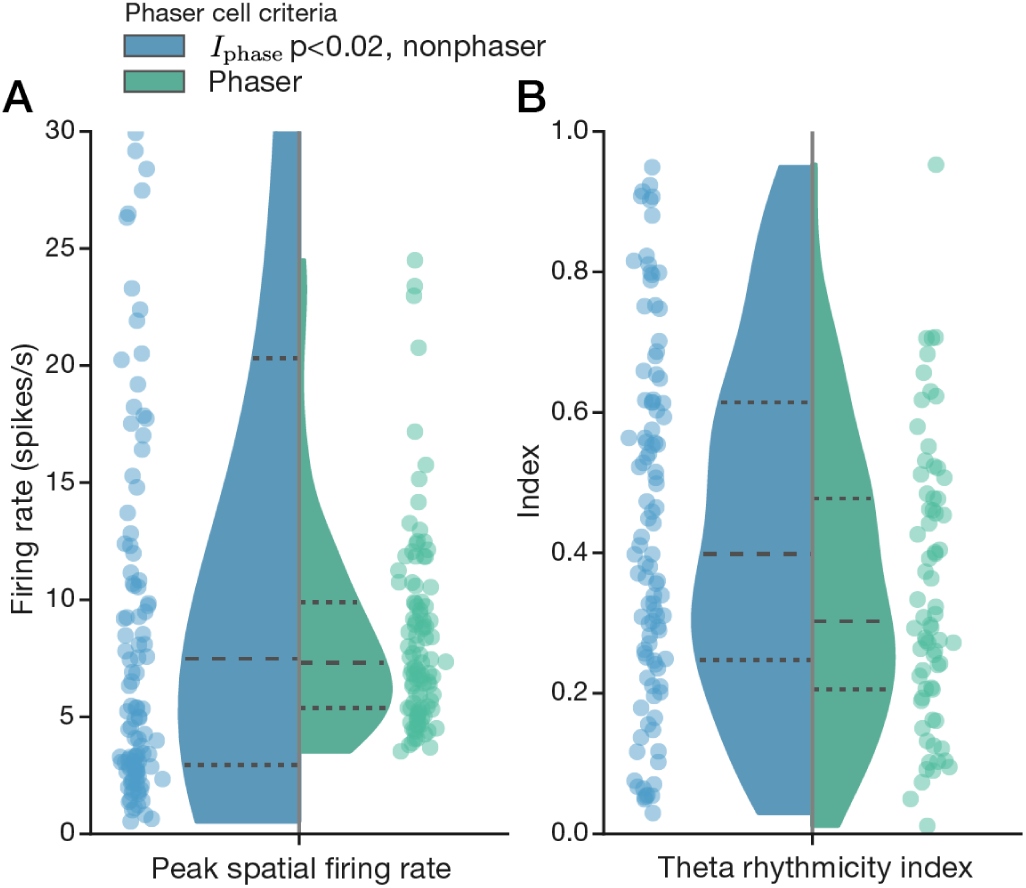
Biological phaser cells have a restricted range of firing rates. The phaser criteria (Results) selected 101/233 recordings from 5/8 rats. Similar to the split distributions in Supplementary Figure 7, we show distributions comparing spatial theta cell recordings not selected (‘nonphaser’; *n* = 233) or selected (‘phaser’; *n* = 101) as biological phaser cells. Peak firing rates (A) for phasers had the same median but a qualitatively restricted range compared to nonphasers (but note that a minimum firing rate of 3.5 spikes/s is one of the phaser criteria and the *y*-axis truncates, for clarity, nonphaser data that is shown in Supplementary Figure 7A). Theta rhythmicity (B) was similarly distributed for nonphaser and phaser recordings.

**Supplementary Figure 10.**
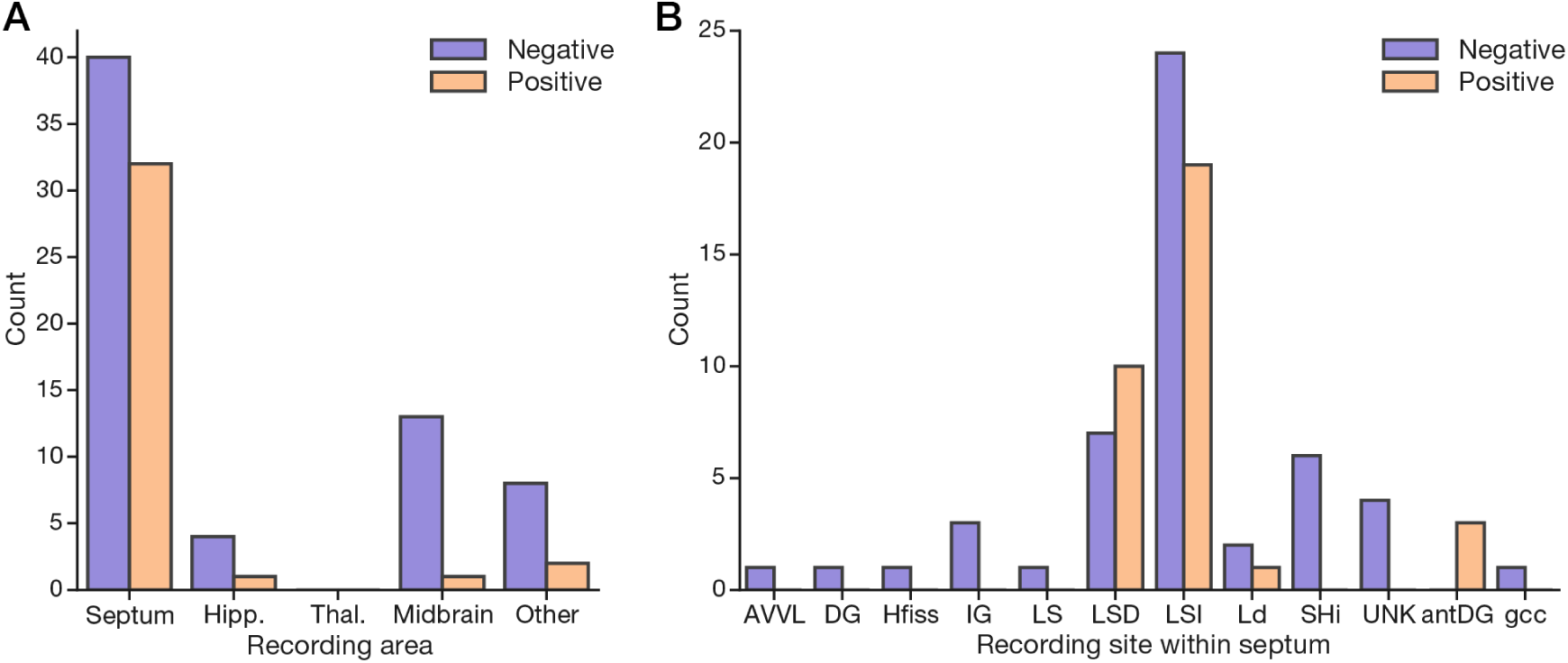
Anatomical distribution of biological phaser cell recordings. (A) Recording counts for brain areas indicating negative and positive biological phasers. Hipp. = hippocampus; Thal. = thalamus; Other includes nucleus accumbens, caudate nucleus, putamen, and CgSHi (TKTK). (B) Recording counts for sites proximal to or within the septum. AVVL = TKTK; (ant)DG = (anterior) dentate gyrus; Hfiss = hippocampal fissure; IG = TKTK; LS(D/I) = lateral septum (dorsal/intermediate); Ld = TKTK; SHi = TKTK; UNK = unknown; gcc = TKTK. **Note: TKTK indicates definitions that will be updated in the next manuscript revision**.

**Supplementary Figure 11.**
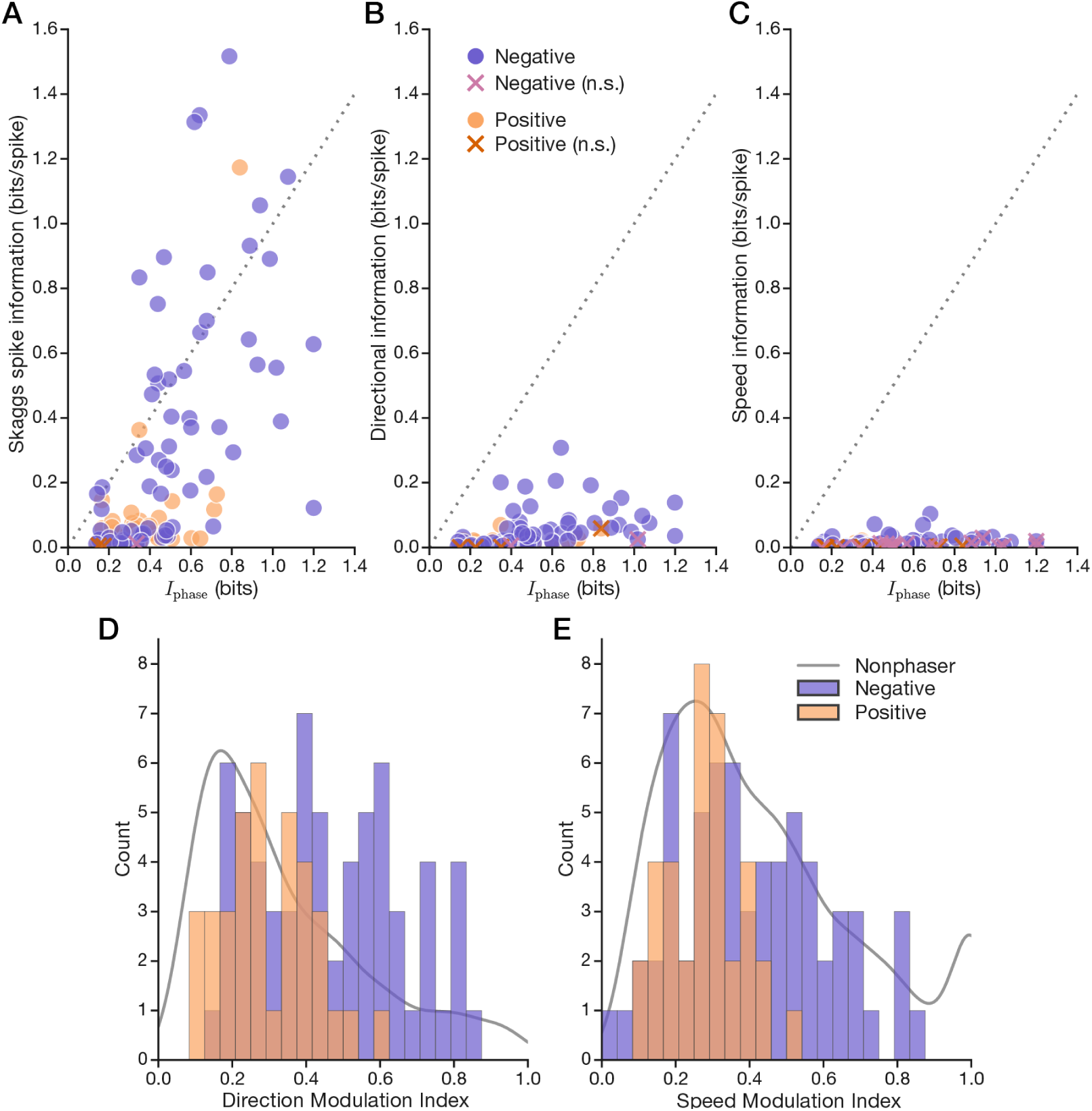
Trajectory modulation reveals bias in biological phaser recordings. The phaser model is based on spatial inputs mediating the correlation between rate and phase. However, biological phasers are recorded in a constrained environment in animals with constrained behaviors, and the hippocampal theta rhythm is strongly speed-modulated (Fuhrmann et al., 2015). (A-C) To evaluate whether the phasers depend on trajectory-based factors, we compare spatial phase information with spike information content (Methods) for position (A, Skaggs measure), direction (B), and speed (C). Most phasers carry strong position information (A), but a minority carry relatively low direction (B) or speed (C) information. (D+E) Histograms of firing-rate modulation indexes for direction (D) and speed (E) for negative/positive phasers (positive composited over negative). Phaser firing rates were substantially modulated by direction and speed. Gray line, kernel-density estimate (0.05 bandwidth Gaussian) of nonphaser recordings (arbitrary scale for comparison).

**Supplementary Figure 12.**
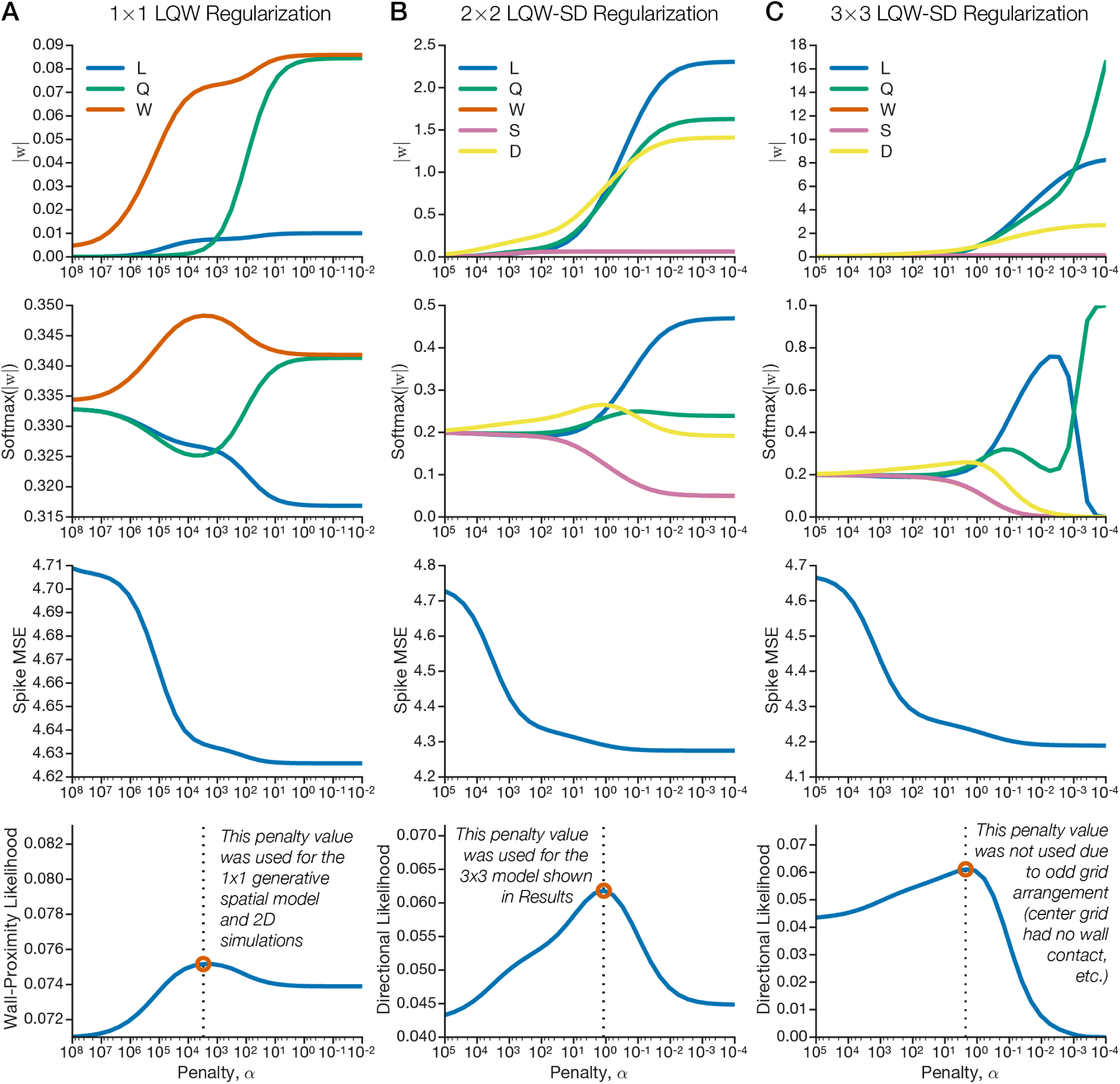
Ridge regularization and shrinkage curves for training GLM models. We trained a series of GLMs to predict spike counts in 300-ms intervals based on spatial (*LQW*) and/or trajectory-based (*SD*) variables (Methods). For the model analysis (Results), the model was trained and tested on a 3 × 3 spatial grid (C); however, the penalty parameter used for training was derived by optimizing the 2 × 2 model (B). Both values were similar, but the 2 × 2 value (B, bottom) was used because the directional likelihood was strongly peaked and the model better captured wall responses (the center grid of the 3 × 3 model was isolated from the walls). The GLM that we used to generate spatial inputs for 2D phaser simulations was only trained on the spatial variables (A, 1 × 1). (top) Absolute model weights for each variable. (second row) Softmax normalization of absolute model weights. (third row) Mean squared error (MSE) of spike-count predictions. (last row) Model likelihood is the the softmax *W* (A) or *D* (B+C) curve divided by the spike MSE (Methods). The maximum likelihood *α* parameter (red circle) is chosen as the *ℓ*_2_-regularization penalty.

**Supplementary Figure 13.**
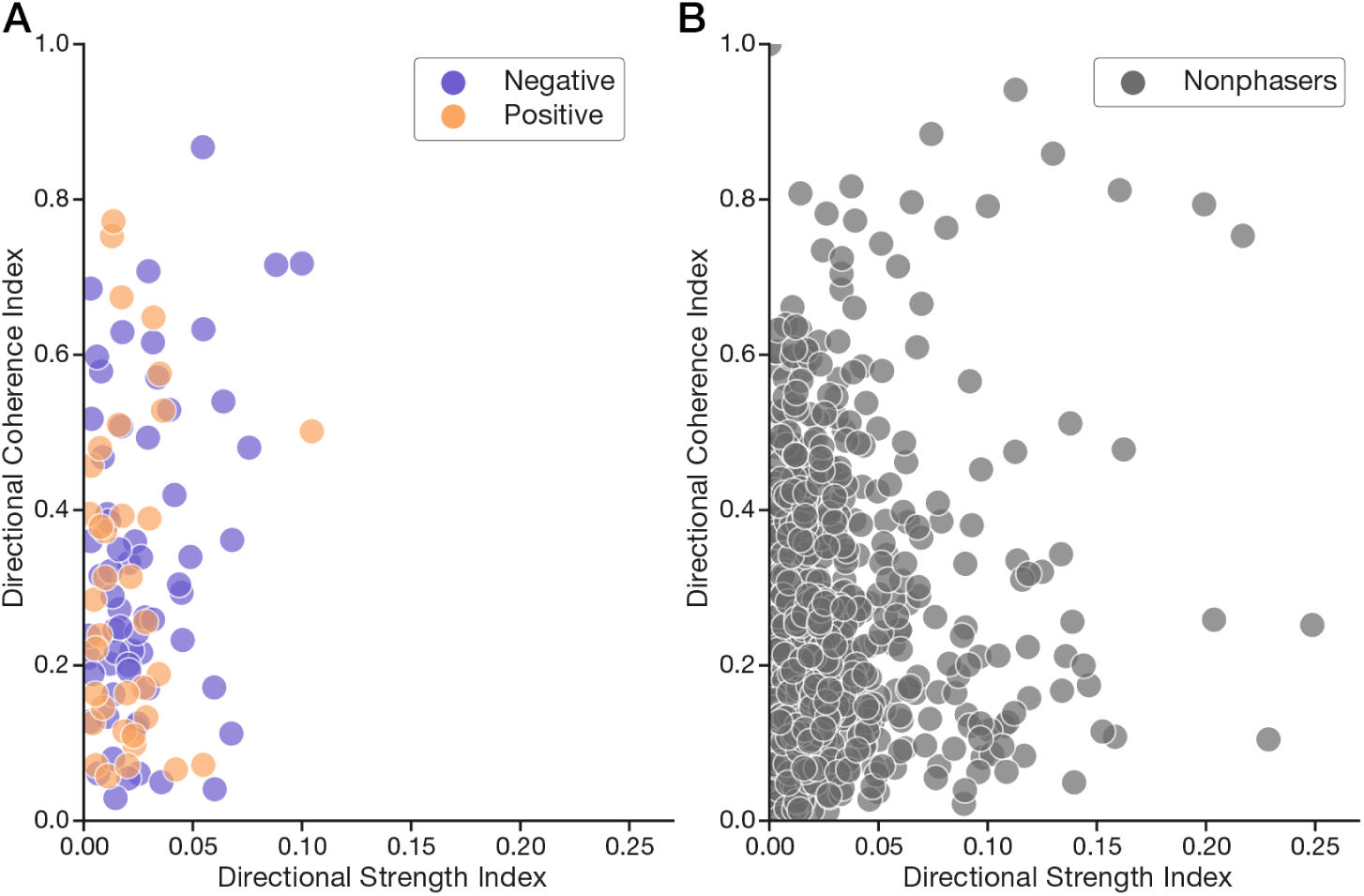
Directional strength and coherence indexes based on GLM weights. The magnitude and direction of the 9 directional vectors from the 3 × 3 GLM were used to compute DSI and DCI, respectively (Methods). Both phasers (A) and nonphasers (B) had a wide range of DCI (*y*-axis), but phasers (A) were restricted to overall low DSI (*x*-axis). The GLM revealed moderately directional responses among nonphasers (B) but only low directionality among phasers (A).

**Supplementary Figure 14.**
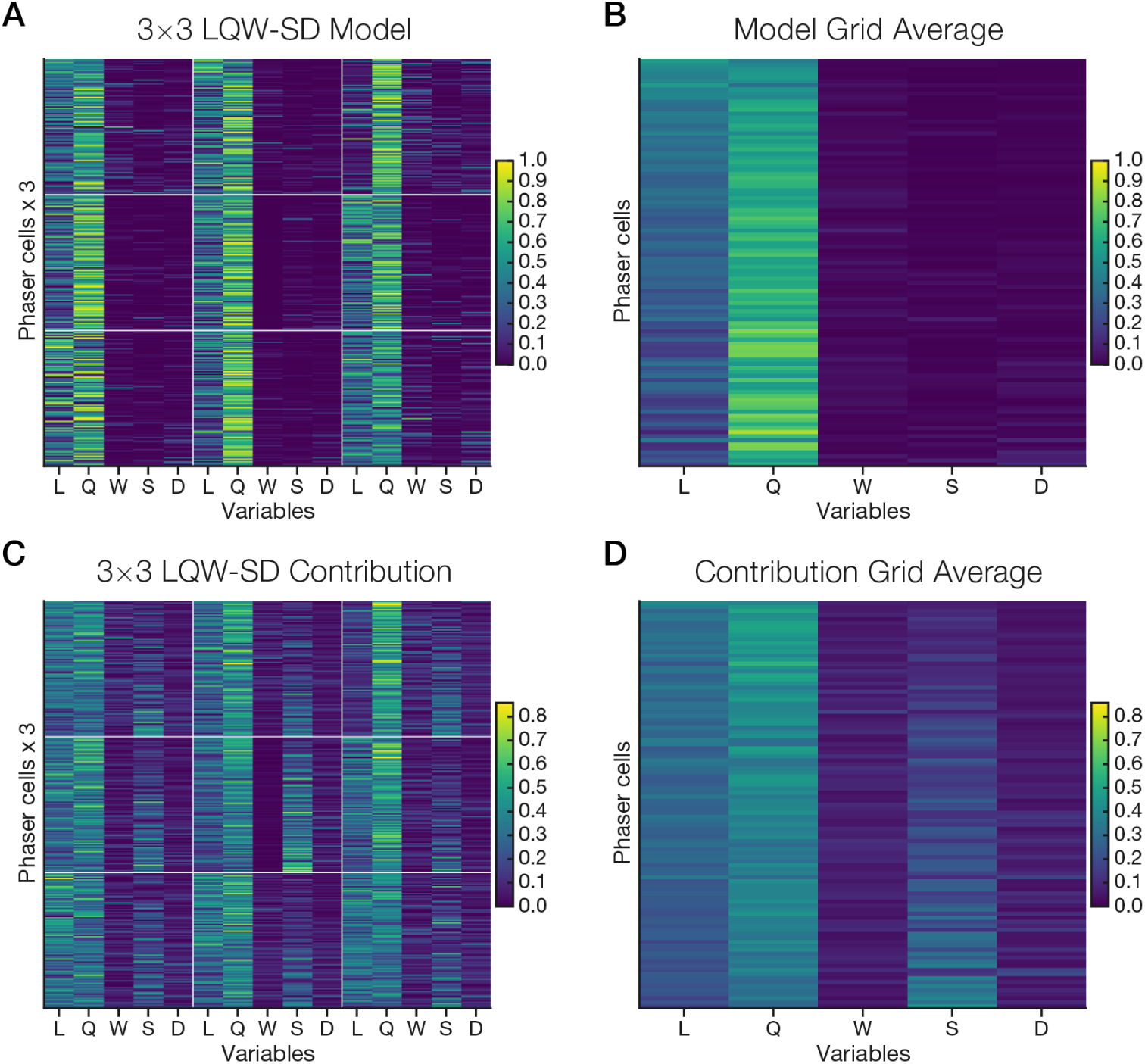
Phaser GLM weights and contributions for every cell and grid section. Full 3 × 3 GLM weights (A+B) and contributions (C+D) for each unique phaser cell (averaged across recordings) are shown in pseudocolor matrix plots. For visualization, cells are presented in the same order in every grid section and grid average, according to the expected value of the cell’s grid-averaged model weights to the left (toward *L*, more spatial) or right (toward *D*, more trajectory-based). To reveal model structure, each variable row in a grid section is sum-normalized and the corresponding grid plots (A+B, C+D) share colorscales.

**Supplementary Figure 15.**
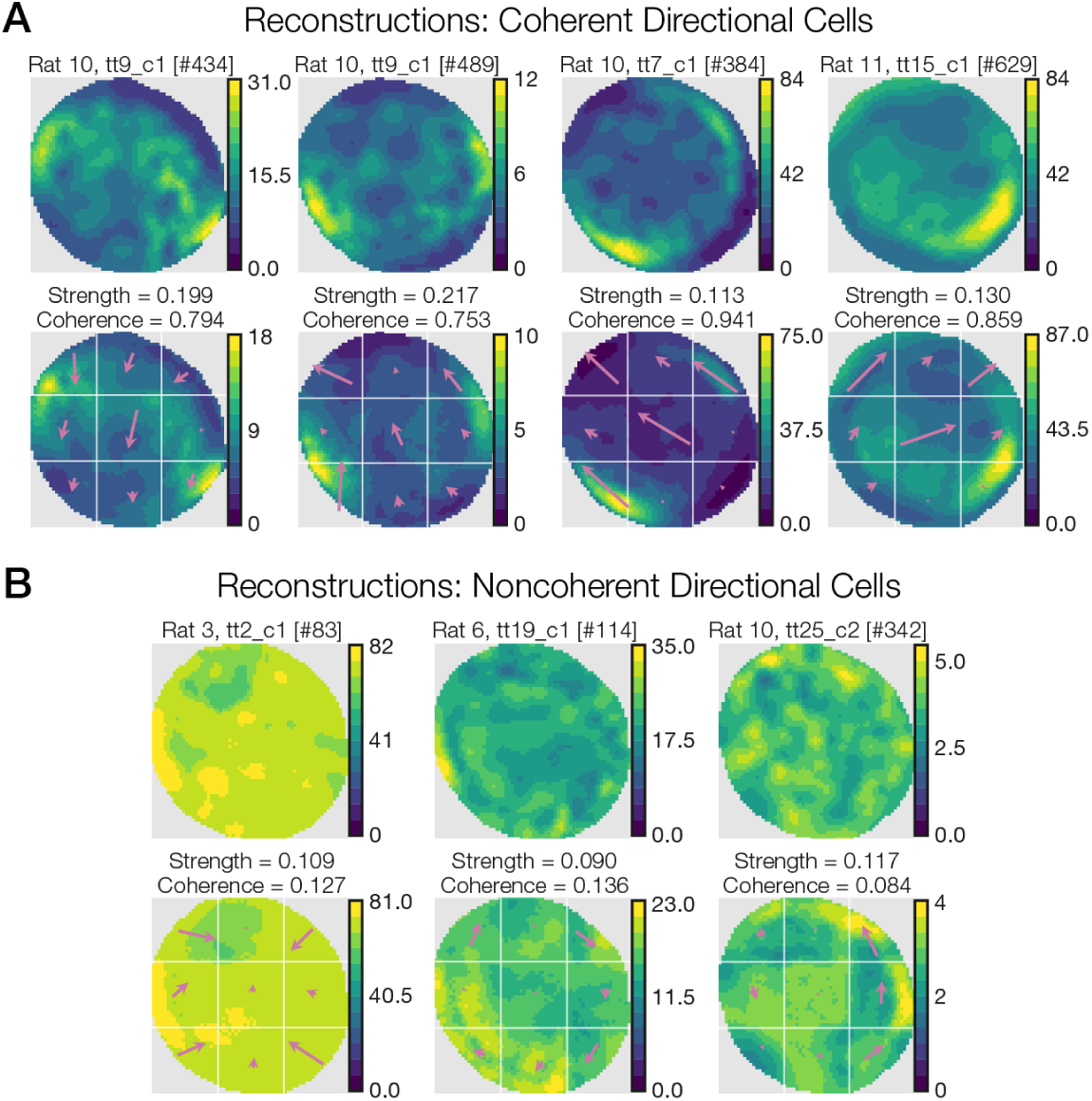
GLM reconstructions of highly directional cells. To show that the LQW-SD 3 × 3 model can accurately reconstruct ratemaps of directional cells, we show examples of coherent (A) and noncoherent (B) directionality. (A) The high peak firing rates and crescent-like spatial modulation indicate that these may be head-direction cells or cells with head-direction inputs. The model’s directional predictors (arrows) are consistently large and well-aligned across grid sections. (B) Recordings with low coherence showed minimal spatial modulation but included directional patterns such as center-facing (left) and clockwise (middle) or anti-clockwise (right) directionality.

**Supplementary Figure 16.**
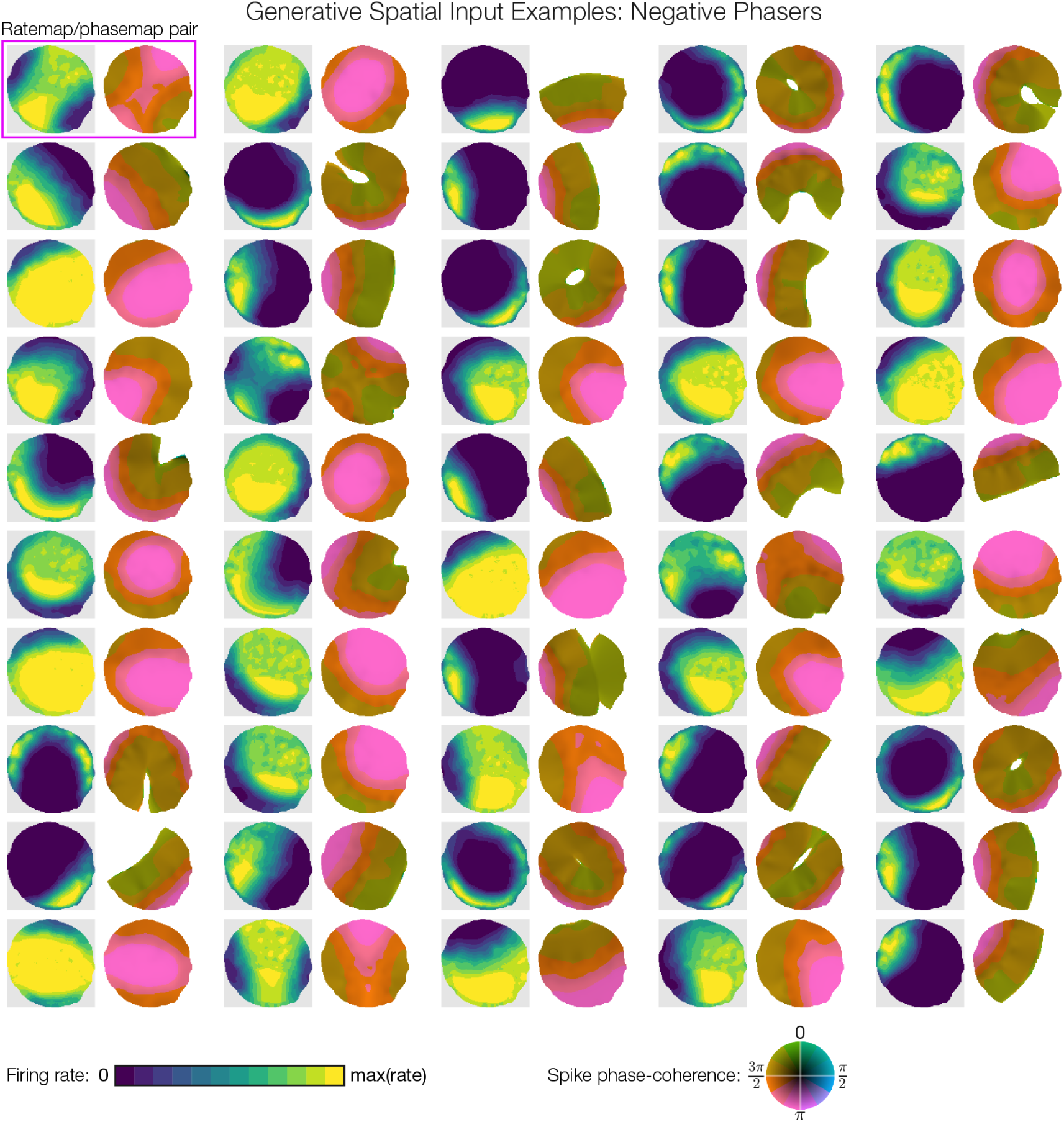
Example negative phasers in 2D synchronization simulations. Ratemap/ phasemap pairs are shown for 50/1,000 negative phasers in the 2D simulations (Figure 9). The rate and phase response of each phaser is driven by a randomly sampled spatial function from the LQW generative model (Figure 8F-J). In the phasemaps, note that the phasers precess from pre-theta-peak (greens; see phase-coherence color wheel at bottom) to theta-trough (pinks) for low- to high-firing regions. Missing phasemap pixels reflect lack of nearby spikes for spatial averaging.

**Supplementary Figure 17.**
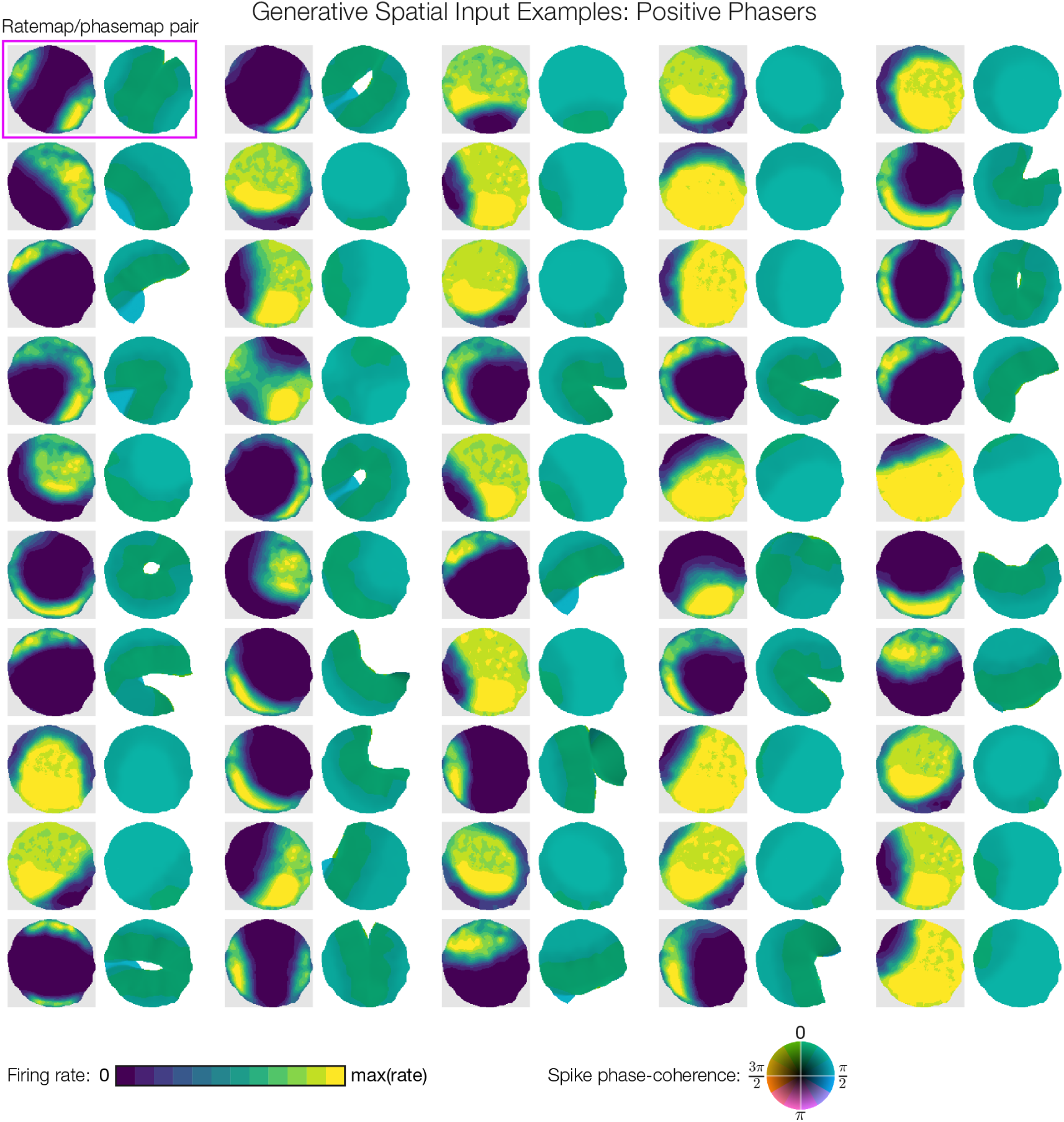
Example positive phasers in 2D synchronization simulations. Ratemap/phasemap pairs are shown for 50/1,000 positive phasers in the 2D simulations (Figure 9). The rate and phase response of each phaser is driven by suppression from a negative phaser with an LQW-generated spatial input (Supplementary Figure 16). In the phasemaps, note that the phasers process from theta-peak (greens) to halfway through the falling phase (blue/green; *π*/2 phase). As for the 1D model (Figure 1) and biological phasers (Figure 6), the positive procession is shallower than the negative precession. Missing phasemap pixels reflect lack of nearby spikes for spatial averaging.

**Supplementary Movie 1.**
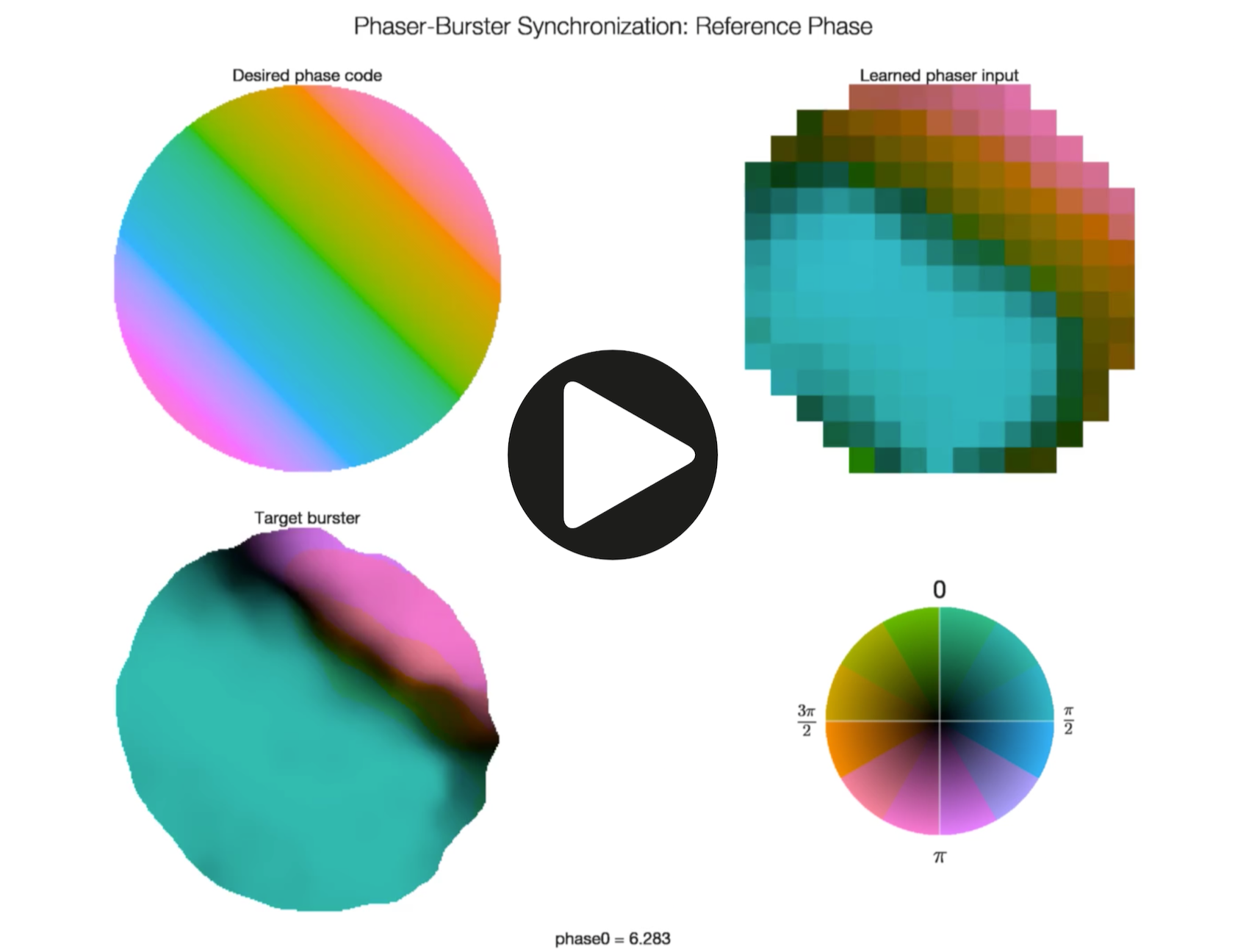
Spatial phaser synchronization across reference phase. The spatial phase codes in Figure 9B differ by reference phase, which determines the spatial offsets of the pattern. Here we show a movie in which the frames iterate through 10-min phaser-target simulations of different reference phases. The desired phase code moves smoothly along the 45° diagonal for a complete cycle so the movie can be looped. The broad negative/positive (pink/blue) synchronization regions compete to cover the environment as the phase code travels.

**Supplementary Movie 2.**
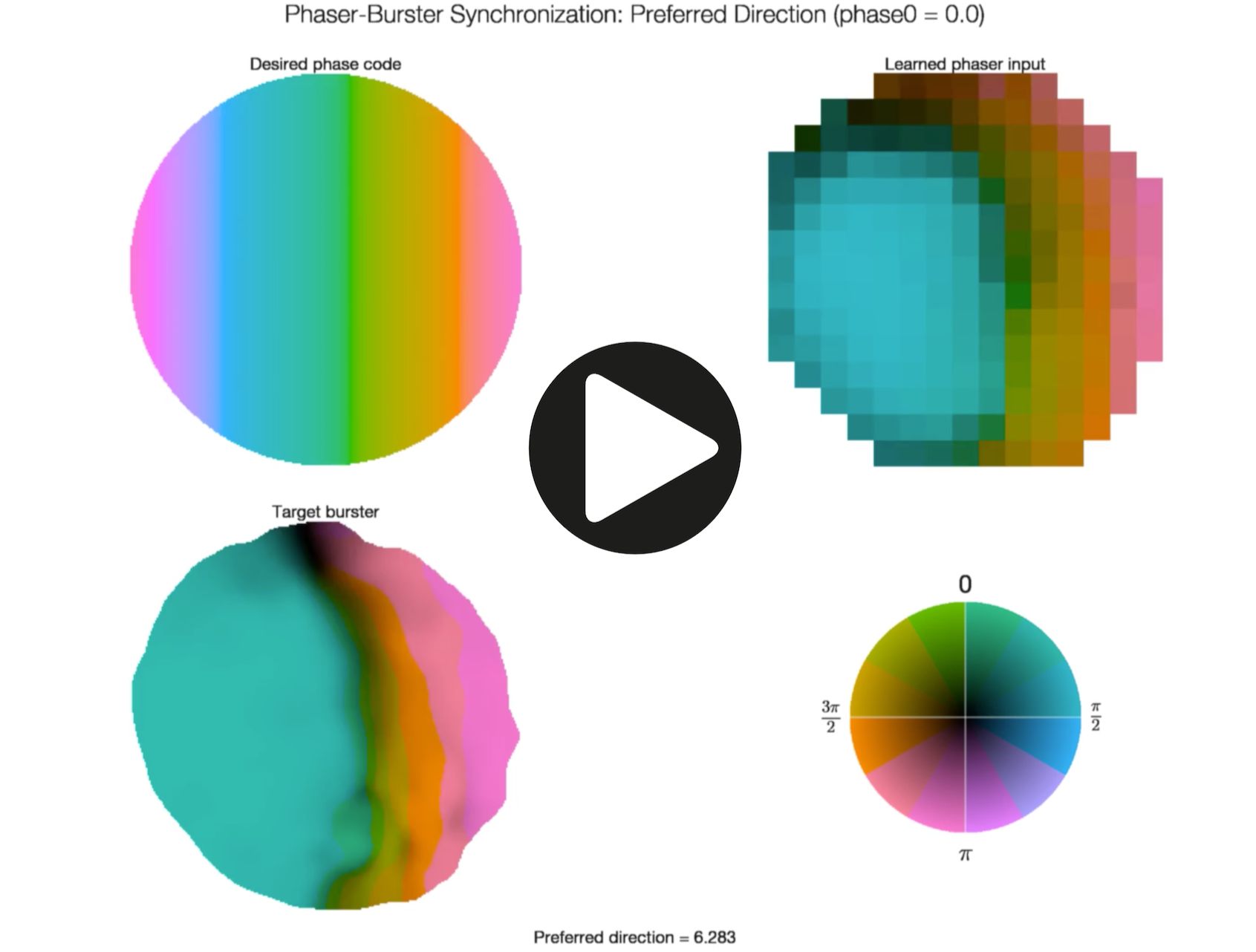
Spatial phaser synchronization across preferred direction: Phase code 1. The spatial phase codes in Figure 9B have a 45° preferred direction, which determines the orientation of the pattern. Here we show a movie in which the frames iterate through 10-min phaser-target simulations of different preferred directions. The desired phase code rotates smoothly for a complete cycle so the movie can be looped. With this reference phase (0.0, at the center of the arena), the negative phasers synchronize a boundary region (oranges/pinks) along the preferred direction as the phase code rotates.

**Supplementary Movie 3.**
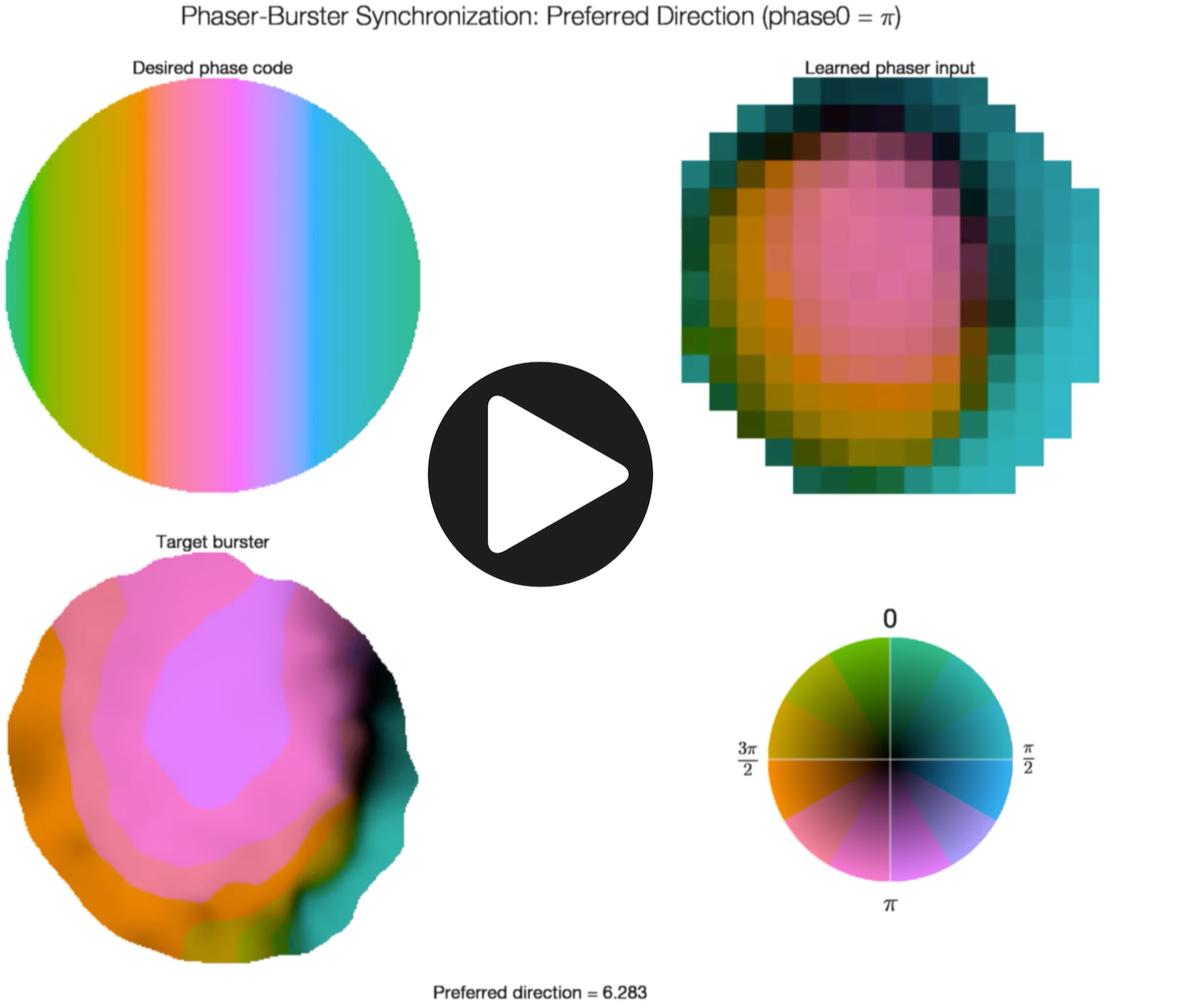
Spatial phaser synchronization across preferred direction: Phase code 2. The spatial phase codes in Figure 9B have a 45° preferred direction, which determines the orientation of the pattern. Here we show a movie in which the frames iterate through 10-min phaser-target simulations of different preferred directions. The desired phase code rotates smoothly for a complete cycle so the movie can be looped. With this reference phase (*π*, at the center of the arena), the positive phasers synchronize a boundary region (blue/green) along the preferred direction as the phase code rotates.

